# Increased levels of the mitochondrial import factor Mia40 prevent the aggregation of polyQ proteins in the cytosol

**DOI:** 10.1101/2021.02.02.429331

**Authors:** Anna M. Schlagowski, Katharina Knöringer, Sandrine Morlot, Ana Sáchez Vicente, Felix Boos, Nabeel Khalid, Sheraz Ahmed, Jana Schramm, Lena Maria Murschall, Per Haberkant, Frank Stein, Jan Riemer, Benedikt Westermann, Ralf J. Braun, Konstanze F. Winklhofer, Gilles Charvin, Johannes M. Herrmann

**Author notes:** to whom correspondence should be sent: Johannes M. Herrmann, Cell Biology, University of Kaiserslautern, Erwin-Schrödinger Strasse 13, 67663 Kaiserslautern, Germany.

## Abstract

The formation of protein aggregates is a hallmark of neurodegenerative diseases. Observations on patient material and model systems demonstrated links between aggregate formation and declining mitochondrial functionality, but the causalities remained unclear. We used yeast as model system to analyze the relevance of mitochondrial processes for the behavior of an aggregation-prone polyQ protein derived from human huntingtin. Induction of Q97-GFP rapidly leads to insoluble cytosolic aggregates and cell death. Although this aggregation impairs mitochondrial respiration only slightly, it interferes with efficient import of mitochondrial precursor proteins. Mutants in the import component Mia40 are hypersensitive to Q97-GFP. Even more surprisingly, Mia40 overexpression strongly suppresses the formation of toxic Q97-GFP aggregates both in yeast and in human cells. Based on these observations, we propose that the posttranslational import into mitochondria competes with aggregation-prone cytosolic proteins for chaperones and proteasome capacity. Owing to its rate-limiting role for mitochondrial protein import, Mia40 acts as a regulatory component in this competition. This role of Mia40 as dynamic regulator in mitochondrial biogenesis can apparently be exploited to stabilize cytosolic proteostasis. (174/175 words)

## Introduction

The accumulation of misfolded or aggregated proteins is frequently observed in stressed cells. Increased levels of misfolded proteins can be the cause, but also the consequence of disease states (for overview see (Chiti & Dobson, 2017, Sontag, Samant et al., 2017, Vaquer-Alicea & Diamond, 2019)). Owing to their capability to sequester chaperones and to occupy proteases, misfolded proteins can route metastable or slowly folding polypeptides into aggregates (Gidalevitz, Ben-Zvi et al., 2006, Kim, Hosp et al., 2016). This might induce a calamitous amplifying reaction which finally leads to proteotoxicity and cell death.

Cells employ a number of different strategies to avoid the hazardous accumulation of misfolded proteins, including their stabilization and disaggregation by chaperones, their degradation by proteases, and their controlled sequestration into aggregates. Mitochondrial functionality modulates cytosolic protein homeostasis in several ways: (1) Protein aggregates often associate with the mitochondrial outer membrane (Gruber, Hornburg et al., 2018, Guo, Sun et al., 2016); in yeast, this property is used to retain misfolded proteins in the mother in order to prevent their inheritance to daughter cells (Böckler, Chelius et al., 2017, Mogk & Bukau, 2014, Zhou, Slaughter et al., 2011). (2) Cytosolic aggregates impair the import of precursor proteins from the cytosol into mitochondria (Cenini, Rub et al., 2016, Li, Vande Velde et al., 2010, Napoli, Wong et al., 2013, Yano, Baranov et al., 2014). (3) *Vice versa*, mitochondrial precursor proteins in the cytosol challenge proteostasis and sequester chaperones and the ubiquitin-proteasome system (Boos, Krämer et al., 2019, Wrobel, Topf et al., 2015). In this context, the hydrophobic carrier proteins of the inner membrane appear to be of specific relevance owing to their hydrophobic nature and specific import mechanism (Wang & Chen, 2015, Williams, Jan et al., 2014). (4) Modulating mitochondrial activities can induce stress resistance pathways and thereby mitigate the accumulation and toxicity of cytosolic aggregates (Fessler, Eckl et al., 2020, Guo, Aviles et al., 2020, Labbadia, Brielmann et al., 2017, Mason, Casu et al., 2013, Sorrentino, Romani et al., 2017). (5) It was proposed that aggregated cytosolic proteins are disentangled on the mitochondrial surface and imported into the mitochondria to enable their degradation by mitochondrial proteases (Li, Xue et al., 2019, Ruan, Zhou et al., 2017) or removal by mitophagy (Guo, Ma et al., 2012, Hwang, Disatnik et al., 2015, Khalil, El Fissi et al., 2015).

The implications of mitochondrial biology for cytosolic proteostasis are even more complicated due to the central role of mitochondria in energy metabolism and redox homeostasis; apparently the pathways relevant for cytosolic aggregate formation and mitochondrial functionality are intertwined in many ways. Yeast cells have been extensively used in the past as a rather simple and well-defined model system to unravel details of cytosolic homeostasis with a special focus on the role of mitochondrial protein import for the formation of cytosolic aggregates (Gruber et al., 2018, Lu, Psakhye et al., 2014, Weidberg & Amon, 2018, Yang, Hao et al., 2016). In order to study the role of mitochondrial protein biogenesis for cytosolic proteostasis, we expressed an established aggregating model protein in the baker’s yeast *Saccharomyces cerevisiae*. The first exon of human huntingtin encodes an aggregation-prone poly-glutamine stretch (polyQ). Expression of polyQ fragments in yeast faithfully recapitulates huntingtin aggregation in a polyQ length-dependent manner (Braun, Buttner et al., 2010). We observed that the expression levels, mitochondrial localization, and functionality of Mia40 are critical determinants for polyQ toxicity. Mia40 (in humans also CHCHD4) is the rate-limiting essential factor of the machinery that imports proteins into the intermembrane space (IMS) of mitochondria (Chacinska, Pfannschmidt et al., 2004, Habich, Salscheider et al., 2019, Naoe, Ohwa et al., 2004, Peleh, Cordat et al., 2016, Peleh, Zannini et al., 2017). Increased levels of Mia40 in the IMS counteract the occurrence of aggregate-inducing nucleation seeds formed by the prion-like protein Rnq1 and suppress the growth arrest induced by an aggregation-prone polyQ protein. Our results show that the regulation of Mia40 levels serves as an efficient molecular mechanism to finetune cytosolic protein homeostasis.

## Results

### Expression of Q97-GFP rapidly induces cytosolic aggregates and stalls cell growth

The first exon of human huntingtin, comprising the polyQ stretch fused to green fluorescent protein (GFP), has been used in the past to study the formation of aggregates in the yeast cytosol (Duennwald, Jagadish et al., 2006, Klaips, Gropp et al., 2020). We expressed a non-aggregating variant of 25 and an aggregation-prone variant of 97 glutamine residues fused to GFP (Q25-GFP and Q97-GFP, Fig. 1A) from a regulatable *GAL1* promoter in wild type yeast cells. Upon induction in galactose-containing medium, Q25-GFP expression resulted in a homogeneous cytosolic distribution whereas Q97-GFP was predominantly forming aggregates seen as punctate signals (Fig. 1B)(Duennwald et al., 2006). Expression of Q97-GFP but not that of Q25-GFP was toxic and prevented cell growth (Fig. 1C).

**Figure 1.**
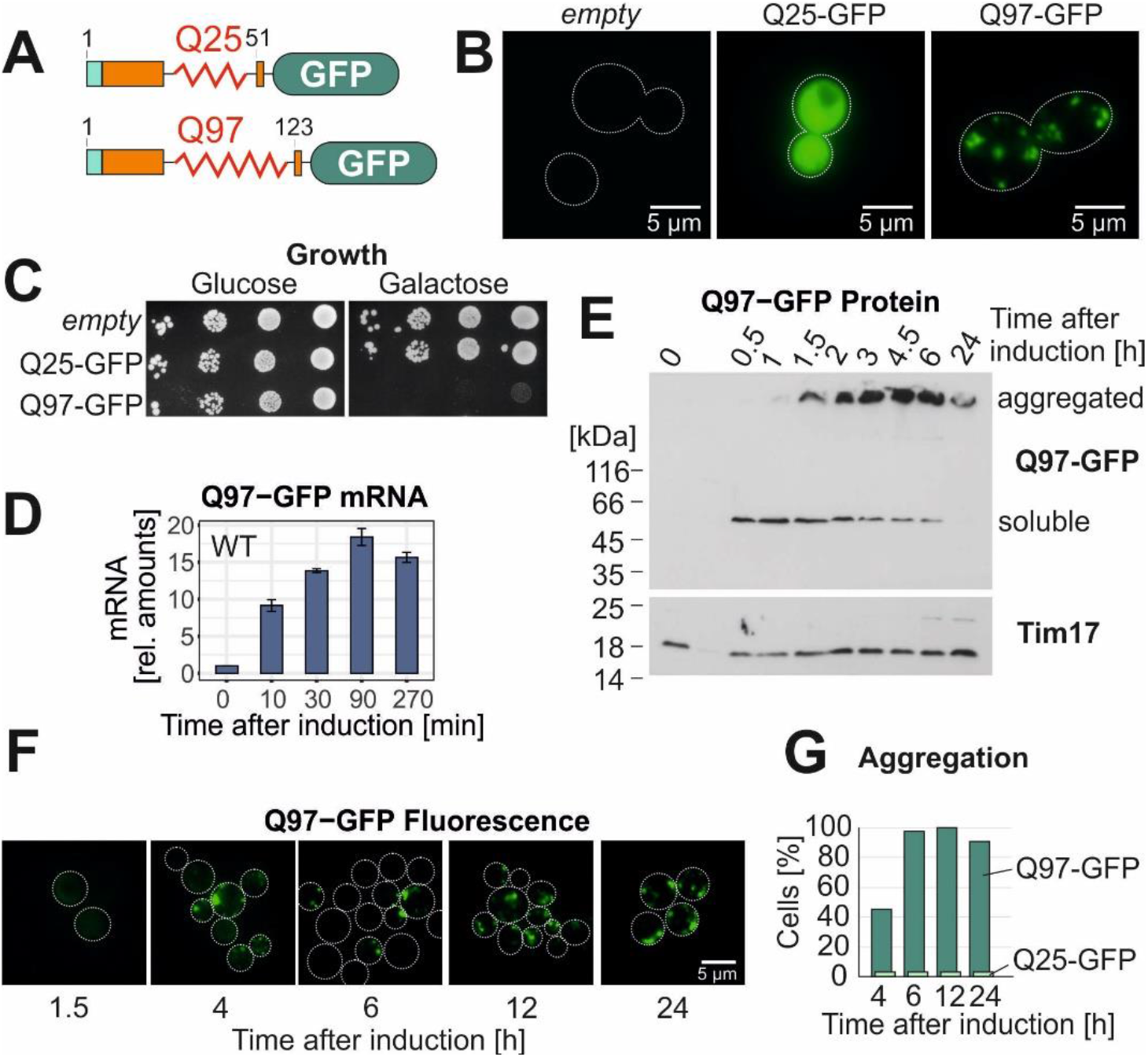
The expression of Q97-GFP in the cytosol leads to the formation of insoluble protein aggregates. **A.** Schematic representation of the structure of the Q25-GFP and Q97-GFP model proteins used in this study. The sequence consists of a FLAG-tag (turquois) followed by the polyQ domain from huntingtin exon1 (orange) and eGFP. **B.** Q25-GFP and Q97-GFP were expressed in wild type cells for 18 h before cells were visualized by widefield fluorescence microscopy. Cells harboring an empty vector are shown for control. **C.** Wild type cells expressing the indicated proteins under control of a galactose-inducible *GAL1* promoter were grown in glucose medium to mid log phase before tenfold serial dilutions were dropped on plates containing glucose or galactose. **D.** Wild type (WT) cells were shifted from glucose to galactose. Then, mRNA levels of Q97-GFP were measured by qPCR. Shown are normalized mean values and standard deviations from three replicates. **E.** Cells were lysed with SDS-containing sample buffer before proteins were visualized by Western blotting. Insoluble aggregates migrate at the top of the gel between stacker and resolving gel. Tim17 served as loading control. **F.** Microscopic images of the Q97-GFP fluorescence. **G.** The percentage of cells containing or lacking detectable aggregates was quantified.

To analyze the order of events in polyQ-mediated toxicity, we analyzed cells after different times of induction. The Q97-GFP mRNA was detectable already after 10 min of induction and reached a maximum after about 90 min (Fig. 1D). The Q97-GFP protein was well observed after 30 min and also reached a maximum after 90-120 min (Fig. 1E). Interestingly, initially the Q97-GFP protein gave rise to a 47 kDa band on SDS gels closely matching the calculated molecular weight of the protein of 42.9 kDa. However, at later time points most of the protein was detected at the upper edge of the gel, indicative for the formation of SDS-insoluble protein aggregates (Douglas, Summers et al., 2009, Kim et al., 2016). This was confirmed by fluorescence microscopy where an initially homogeneous GFP signal was replaced by aggregates several hours after induction in an increasing number of cells (Fig. 1F). After 4 h, about 50% of all cells that showed green fluorescence contained aggregates which increased to almost 100% after 6 h of expression (Fig. 1G). Thus, the Q97-GFP protein is initially soluble and homogeneously distributed in the cytosol, but then rapidly aggregates in basically all cells, which coincides with a growth arrest.

### A functionally compromised mutant of the mitochondrial import factor Mia40 shows increased Q97 toxicity

Previous studies suggested that cytosolic protein aggregation can disturb the functionality of mitochondria (Mossmann, Vogtle et al., 2014, Papsdorf, Kaiser et al., 2015, Solans, Zambrano et al., 2006). We therefore tested whether the Q97-GFP aggregates colocalize with mitochondria (Fig. 2A). We observed that mitochondria and aggregates were clearly distinct structures. Mitochondrial colocalization that was seen for a few puncta might represent random contacts. Moreover, the formation of the Q97-GFP aggregates did not destroy the mitochondrial network which maintained its reticular structure for at least 4 h after induction of Q97-GFP expression (Fig. 2A). We also monitored the respiration-driven oxygen consumption and the activity of cytochrome *c* oxidase which was only moderately reduced, even after 24 h of Q97-GFP expression (Fig. 2B, C). We conclude that the expression of Q97-GFP induces the formation of cytosolic aggregates and impairs cell growth, however, at least within the first hours of induction, it only has mild effects on mitochondrial morphology and the functionality of the respiratory chain.

**Figure 2.**
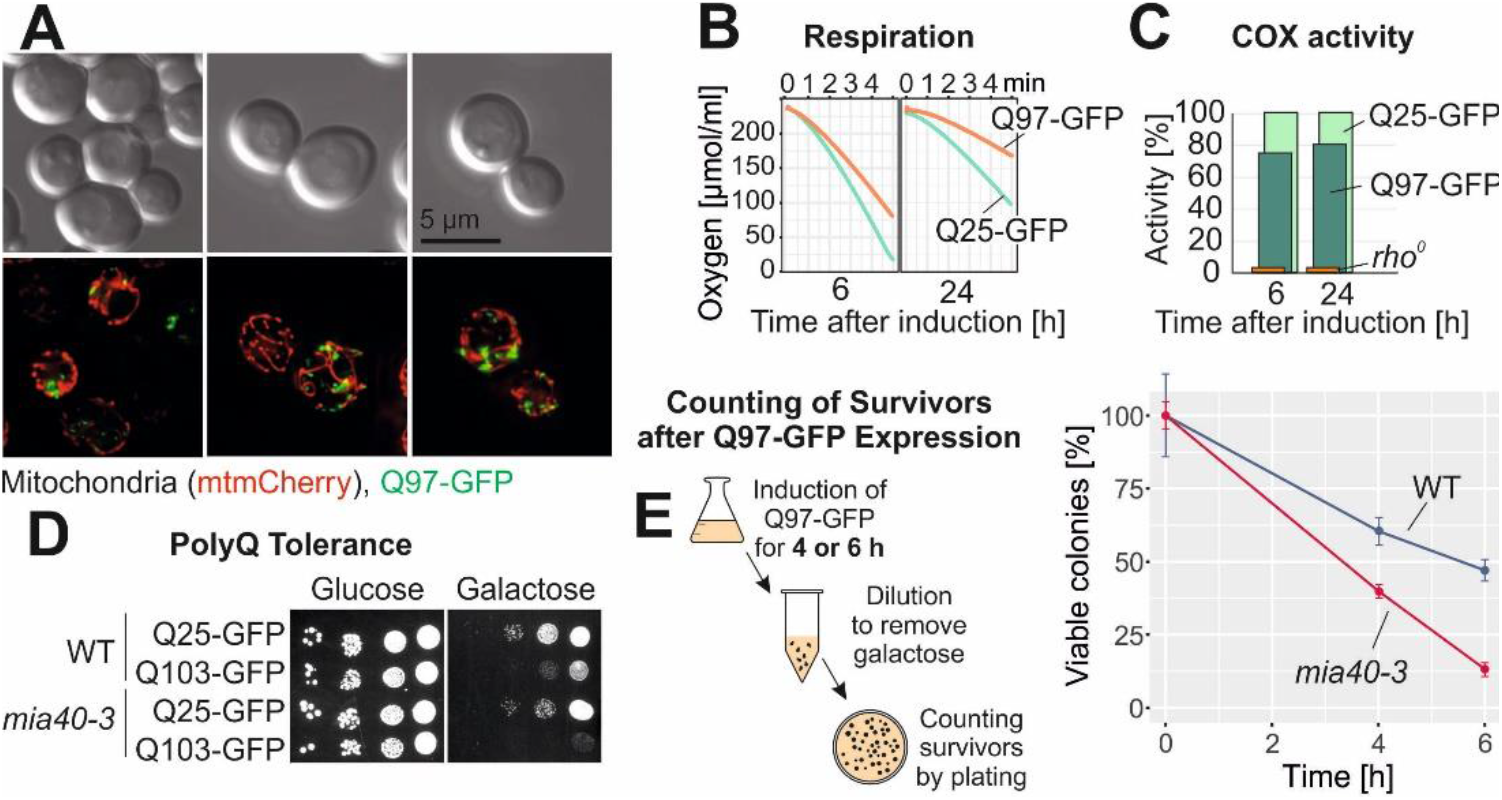
Temperature-sensitive *mia40-3* cells are hypersensitive to polyQ aggregates. **A.** Mitochondria were visualized by mitochondrially targeted mCherry (red) in WT cells expressing Q97-GFP for 4 h. Fluorescence micrographs are maximum intensity projections of z stacks subjected to deconvolution. **B, C.** The polyQ proteins were expressed for the indicated time periods. Mitochondria were isolated and their ability to respire (shown as NADH-induced oxygen consumption) or their activity of cytochrome *c* oxidase (shown as their capacity to oxidize reduced cytochrome *c*) was measured. Mean values of three replicates are shown. **D.** Cells of the temperature-sensitive *mia40-3* mutant (Chacinska et al., 2004) or the corresponding wild type were transformed with plasmids expressing the indicated proteins from a low-expression *GALL* promoter (Mason et al., 2013) and grown on the respective carbon sources at permissive conditions (25°C). **E.** Q97-GFP was expressed in *mia40-3* and corresponding wild type cells for the times indicated. Cells were harvested, washed and survivors were counted after plating on glucose plates. Mean values and standard deviations of three replicates are shown.

Next, we tested whether mitochondrial functionality influenced the polyQ toxicity. Since the expression of the Q97-GFP from the *GAL* promoter was highly toxic, we used a plasmid which expressed Q103-GFP from a low-expression *GALL* promoter (Mason et al., 2013) and tested its effect on the growth in a temperature-sensitive Mia40 (*mia40-3*) mutant. Mia40 is an essential protein of the mitochondrial protein import machinery (Chacinska et al., 2004, Naoe et al., 2004) and the discovery that cytosolic precursors of mitochondrial proteins are toxic was initially made in such a temperature-sensitive Mia40 mutant (Wrobel et al., 2015). We observed that the *mia40-3* mutant was more sensitive to Q103-GFP expression than the wild type: even at permissive temperatures cells were unable to grow on galactose-containing plates (Fig. 2D). This suggests that the burden of non-imported mitochondrial precursors in the cytosol adds to the problems caused by toxic polyQ proteins.

To test whether the effects on growth are due to a growth arrest or to cell death, we expressed high levels of Q97-GFP in wild type and *mia40-3* cells and grew them to log phase. Cells were exposed for 4 or 6 h to galactose, reisolated and washed before surviving cells were counted by a plating assay on glucose medium. We observed that viability of Q97-GFP expressing cells rapidly declined, in particular in the *mia40-3* mutant (Fig. 2E), suggesting that the growth arrest is caused by cell death (Chacinska, Lind et al., 2005).

### Overexpression of Mia40 suppresses the aggregation and toxicity of Q97-GFP

The observed hypersensitivity of *mia40-3* cells to Q97-GFP expression inspired us to test the effect of Mia40 overexpression. To this end, we made use of a strain which expresses Mia40 from a *GAL* promoter and which, in addition, harbors a plasmid with *MIA40* under control of its endogenous promoter (Terziyska, Grumbt et al., 2007). This extra-copy of *MIA40* allows the strain to grow on glucose where the *GAL*-driven expression is repressed. This strain was transformed with the Q25-GFP and Q97-GFP expression plasmids. To our surprise, we observed that co-overexpression of Mia40 indeed strongly protected cells against the Q97-GFP-mediated growth arrest (Fig. 3A). The Mia40-induced suppression was not due to reduced Q97-GFP expression as *GAL*-Mia40 cells even contained higher Q97-GFP levels than wild type cells (Fig. 3B). Intriguingly, upon Mia40 overexpression Q97-GFP hardly formed any SDS-resistant aggregates, indicating that elevated levels of Mia40 suppressed aggregation of Q97-GFP.

**Figure 3.**
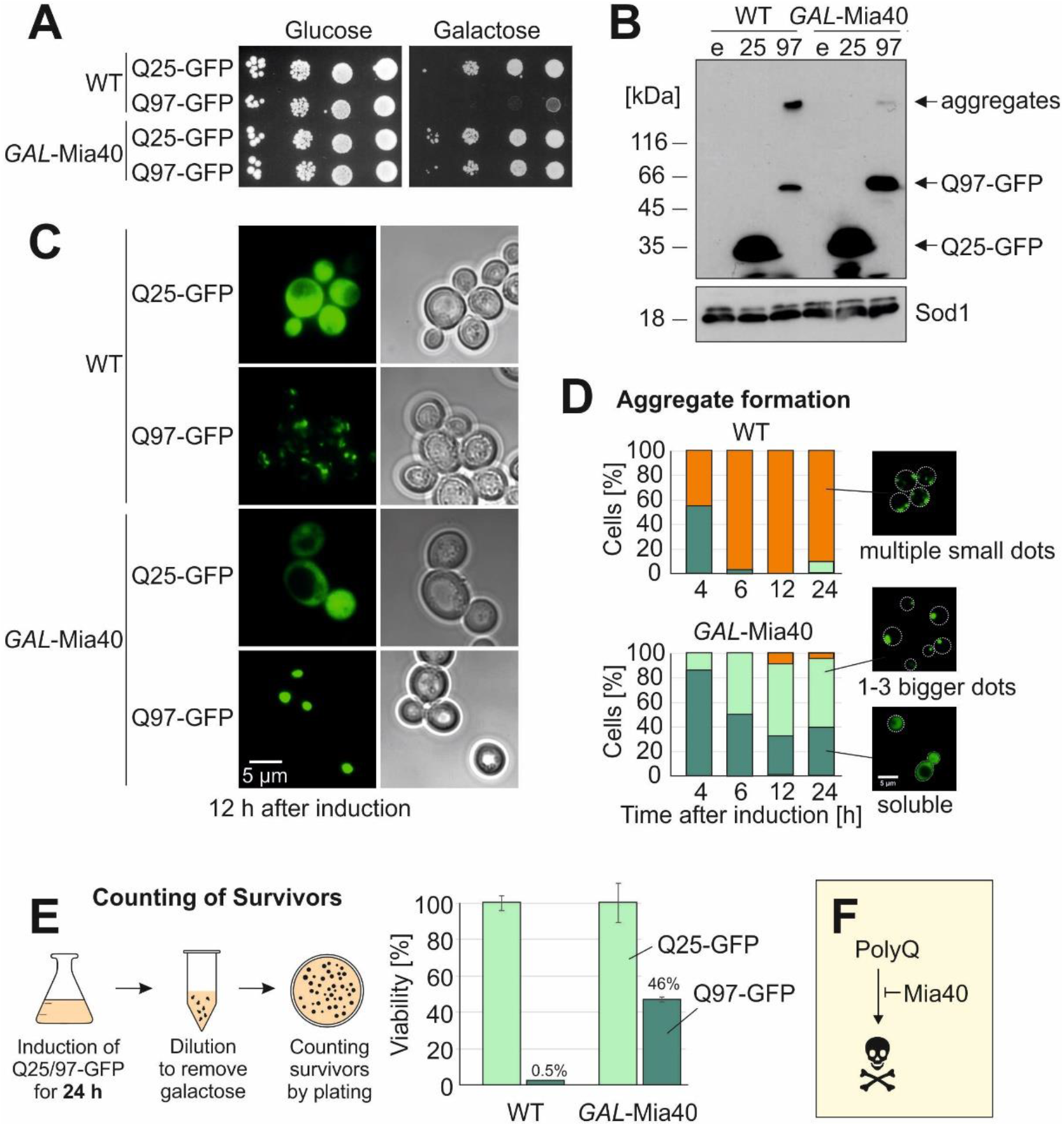
Overexpression of Mia40 suppresses polyQ toxicity in yeast. **A.** Indicated strains were grown on glucose medium before tenfold serial dilutions were dropped on glucose- or galactose-containing plates. **B.** Cell extracts were analyzed by Western blotting after shifting cultures to galactose for 16 h. **C.** Microscopy images of the indicated strains 12 h after shifting them to galactose. Note that in *GAL*-Mia40 cells the form and number of aggregates is very different to WT cells. **D.** The patterns of the Q97-GFP distribution were quantified after different time of expression, n=100. **E.** PolyQ proteins were expressed for 24 h before survivors were counted. Mean values and standard deviations from three independent experiments are shown. **F.** The *GAL*-Mia40 strain was much more resistant to Q97-GFP expression than WT cells, indicating that high Mia40 levels can suppress polyQ toxicity.

Fluorescence microscopy confirmed the striking difference in aggregate formation (Fig. 3C, D): whereas Q97-GFP formed several small, distinct aggregates in wild type cells, the protein was either homogeneously dispersed in the *GAL*-Mia40 overexpression strain or formed one large intracellular aggregate. Such large polyQ aggregates were described before in mutants of cytosolic chaperones (Dehay & Bertolotti, 2006, Higgins, Kabbaj et al., 2018, Krobitsch & Lindquist, 2000, Meriin, Zhang et al., 2002, Yang et al., 2016).

Upon expression of Q25-GFP or Q97-GFP, mitochondria still formed a wild type-like reticulate structure, both in the presence and absence of Mia40 overexpression (Figure EV1). In some cells that simultaneously overexpressed Q97-GFP and Mia40 we noticed more reticulate, ‘curly’ mitochondria (Figure EV1). These structures were not observed when only one of these proteins was overexpressed. We are not aware that such structures were reported before, and their significance is currently unclear.

Mia40 overexpression was very effective in the repression of polyQ toxicity and about half of all cells even survived the *GAL*-driven expression of Q97-GFP for 24 h (Fig. 3E). From this we conclude that the overexpression of the mitochondrial import factor Mia40 can protect against the formation of toxic polyQ aggregates (Fig. 3F).

### Mia40 influences the formation and inheritance of polyQ aggregates

Since we had observed that induction of Q97-GFP in wild type cells resulted in an initially homogeneously distributed protein that later formed SDS-resistant aggregates, we wanted to better understand the temporal and spatial patterns of aggregate formation and inheritance in wild type and *GAL*-Mia40 cells. To this end, we monitored the growth of wild type and *GAL*-Mia40 cells after galactose-driven induction of Q25-GFP and Q97-GFP in a microfluidics growth chamber (Morlot, Song et al., 2019) under the fluorescence microscope (Fig. 4A). Thereby, individual cells could be followed over time which showed that *GAL*-Q25 levels did not interfere with bud formation and growth. In contrast, in wild type cells Q97-GFP expression slowed down cell division, and cells died prematurely (Fig. 4B, Supplemental Movies 1 to 3). The pattern was entirely different in *GAL*-Mia40 cells: Cells accumulated Q97-GFP with a homogeneous intracellular distribution and thereby reached higher fluorescence levels than wild type cells (Fig. 4B, C). However, we noticed that, in individual cells, occasionally all the evenly distributed fluorescence collapsed indicating the formation of one characteristic large aggregate (Fig. 4B). This collapse was very sudden and occurred from one frame to the next, i.e. in less than 10 min and without changing the total fluorescence intensity of a cell (Fig. 4D). There was no noticeable correlation between the level of Q97-GFP signal and the time point of the collapse, suggesting that the formation of these large aggregates was a stochastic process. Once cells had these large aggregates formed, they ceased to produce new buds. Obviously, they were arrested in cell cycle and died after some time (Supplemental Movie 4). Even when galactose was removed to stop Q97-GFP expression, these aggregates persisted whereas the diffuse Q97-GFP was degraded (Fig. 4E).

**Figure 4.**
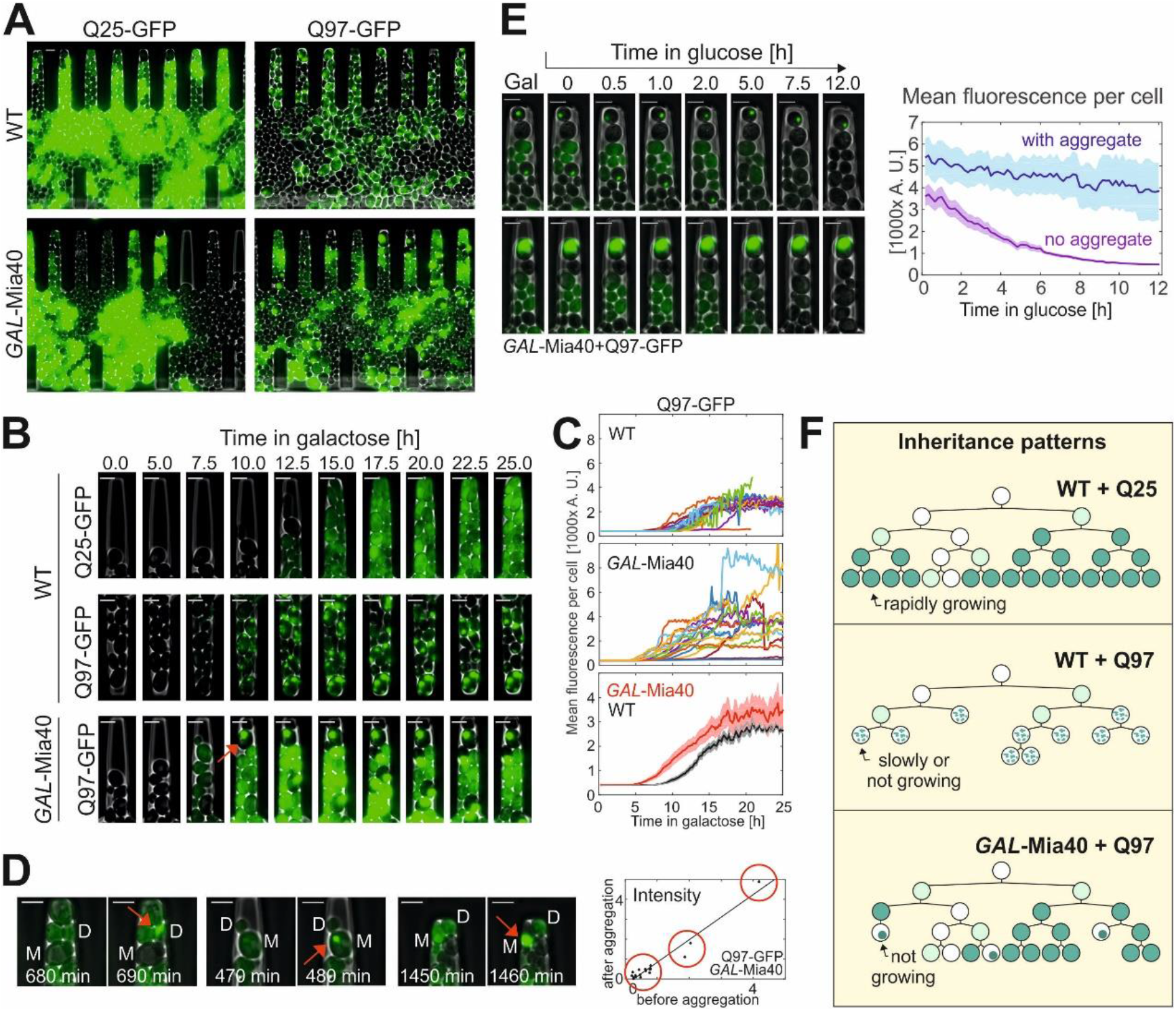
Q97-GFP shows a sporadic aggregation behavior in *GAL*-Mia40 cells. **A.** Cells were grown in microfluidic chambers (see supplemental Videos 1-4). To better visualize the fluorescence signals different intensity settings were used, so that the intensities here are not comparable. **B.** Still frames after different times of growth indicate a uniform fluorescence in Q25-GFP-expressing cells. Q97-GFP expression in wild type leads to many small and scattered aggregates per cell. In *GAL*-Mia40 cells, Q97-GFP accumulates to much larger intensities but then suddenly collapses into one single aggregate per cell. The arrow depicts aggregate formation. **C.** Quantified signal intensities in cells of the indicated mutants. In top and middle graphs, each curve corresponds to a single cell. Averaged signals (mean + SEM) in bottom graph (N=20 for WT, N=17 cells for *GAL*-Mia40). **D.** Still frames directly before and after the Q97-GFP aggregate formation in *GAL*-Mia40 cells. Total cellular fluorescence intensity before aggregation and total intensity in the aggregate were quantified and are shown in the graph on the right hand side, indicating that the entire cellular fluorescent signal collapses into the one single aggregate. **E.** Cells were shifted from galactose to glucose to stop expression of Q97-GFP. The quantified image shows mean values and SEM from 20 cells with aggregates and 15 cells without aggregate, indicating that soluble Q97-GFP is degraded whereas the aggregated protein is of much higher stability. **F.** Schematic inheritance patterns of polyQ proteins in wild type and *GAL*-Mia40 cells. Compare also Fig. EV2 for more details.

To better follow the distribution of the GFP patterns in these cells we developed an artificial intelligence tool which was trained to recognize individual yeast cells and the different patterns within these cells (i.e. unstained, homogeneous, smaller aggregates, one large aggregate) (Fig. EV2A). This algorithm quantified in an unbiased manner the different categories of cells in the four strains (Fig. EV2B). These numbers allowed us to distinguish three distinct scenarios (Fig. 4F): (1) soluble Q25-GFP homogeneously accumulates in all cells over time. (2) Q97-GFP forms small aggregates in basically all wild type cells, slows down growth and leads to premature cell death. (3) In *GAL*-Mia40 cells several different patterns emerged: some cells formed the large aggregates which stopped their growth, whereas other cells continued to grow and accumulated homogeneously distributed Q97-GFP. Thus, the large aggregates of the *GAL*-Mia40 cells are not better tolerated than the aggregates in the wild type, but seem to be even more toxic. However, since they occur only sporadically, cells forming these structures are outnumbered by well growing cells showing homogeneous Q97-GFP staining.

### Mia40 overexpression reduces the number of cells with Rnq1 aggregation seeds

What is the molecular basis of the different aggregation phenotypes in wild type and *GAL*-Mia40 cells? To address this, we induced Q97-GFP expression for 4.5 h in both strains, lysed the cells with the non-denaturing detergent NP-40 and subjected the NP-40-soluble (non-aggregated) fraction to proteomics by quantitative mass spectrometry (Fig. 5A). This revealed, as expected, increased levels of Mia40 in the *GAL*-Mia40 strain (about 6 times more than in cells with normal Mia40 levels) and slightly increased levels of a number of mitochondrial proteins, including some Mia40-dependent IMS proteins (Extended View Table EV2) confirming previous studies (Peleh et al., 2016, Wrobel et al., 2015). Interestingly, the NP-40-soluble fractions from the *GAL*-Mia40 strain showed altered levels of Rnq1 which is a well-characterized yeast prion protein that serves as crystallization seed in the aggregation process of many proteins, including polyQ proteins (Douglas et al., 2009, Duennwald et al., 2006, Meriin et al., 2002, Wolfe, Ren et al., 2013).

**Figure 5.**
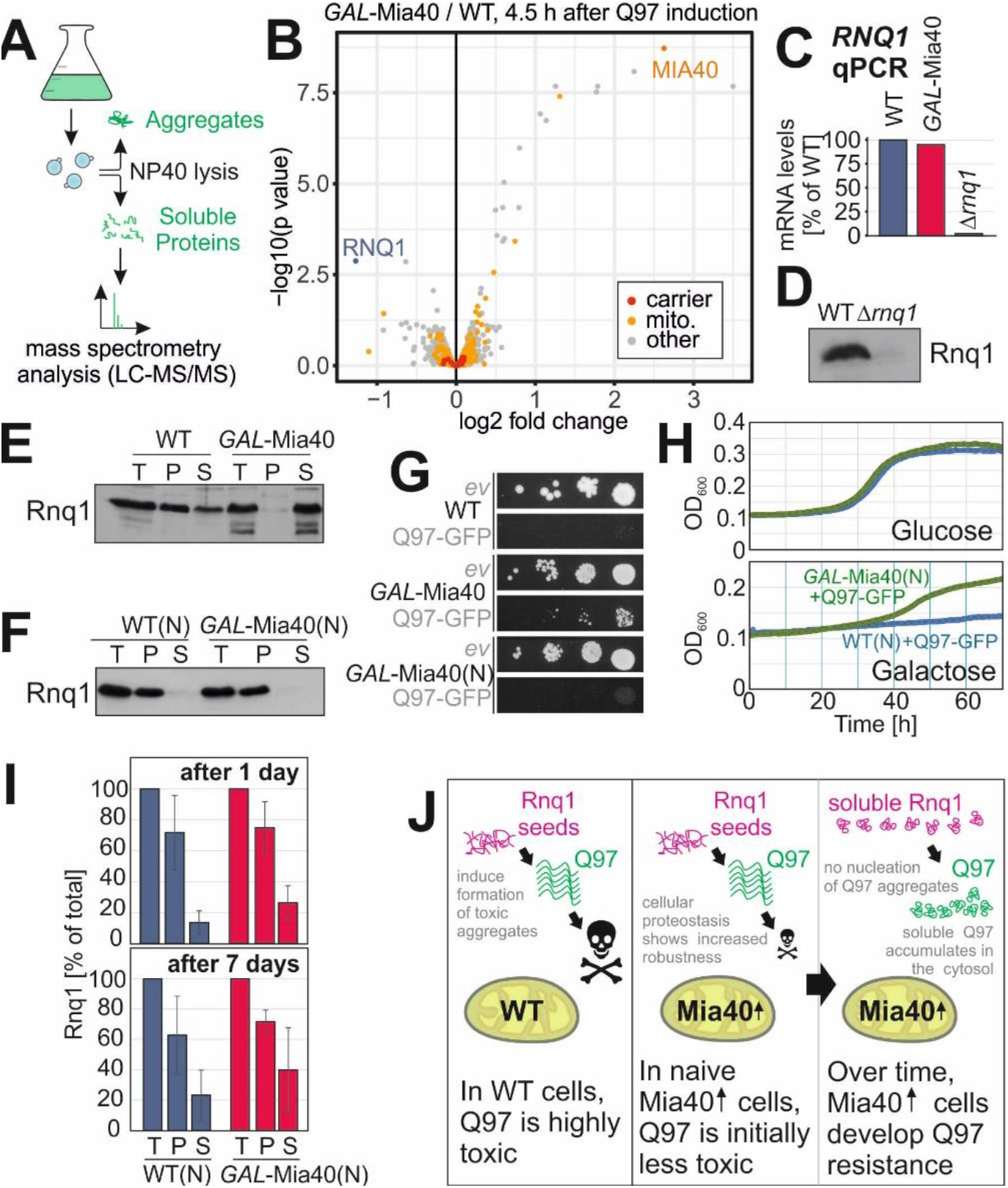
Overexpression of Mia40 leads to increased proteostatic robustness of the cytosol. **A.** Workflow of the quantification of soluble proteins by mass spectrometry. **B.** Q97-GFP was expressed for 4.5 h in wild type and *GAL*-Mia40 cells in two biological replicates each. Cells were lysed with NP-40 before soluble proteins were quantified. Mitochondrial proteins (mito.) and members of the mitochondrial carrier family (carrier) are visualized in a volcano plot. P-values were derived from a moderated t-test (limma). **C.** Transcript levels for *RNQ1* were measured by qPCR in the strains indicated. **D.** Antibodies against Rnq1 were used in Western blots to specifically detect the protein in whole cell extracts of yeast cells. **E, F.** Cell extracts of the strains indicated were separated into soluble (S) and aggregated (P) fractions. Rnq1 was detected by Western blotting. Whole cell extracts (total, T) were loaded for control. The wild type and *GAL*-Mia40(N) strain shown in F were newly generated. **G.** Drop dilution experiments of the strains indicated on galactose plates; *ev*, empty vector. **H.** Growth curves of the indicated strains at 30°C. **I.** Cells of the newly made strains were grown on galactose for 1 or 7 days, respectively. Cells were lysed before the distribution of Rnq1 in the soluble and pellet fractions was quantified. Shown are mean values and standard deviations of three independent replicates. **J.** Mia40 overexpression reduces the toxicity of polyQ proteins. In newly made cells, Q97-GFP is still toxic (though less than in wild type) but after several generations, cells escape sensitivity by loss of Rnq1 nucleation seeds.

The transcription of *RNQ1* was not altered in *GAL*-Mia40 cells as judged from qPCR signals (Fig. 5C). We therefore raised Rnq1 antibodies which showed comparable Rnq1 protein levels in both strains. However, a large fraction of Rnq1 formed aggregates in wild type cells, whereas Rnq1 was soluble in the *GAL*-Mia40 strain (Fig. 5D, E). The absence of Rnq1 seeds in *GAL*-Mia40 cells might explain the alteration in Q97-GFP aggregation. Indeed, *Δrnq1* cells showed a comparable or even more pronounced Q97-GFP resistance than the *GAL*-Mia40 mutant (Fig. EV3A).

To test whether the increased Mia40 levels were causative for the changed Rnq1 aggregation, we generated newly made versions of the wild type and *GAL*-Mia40 mutants (Fig. 5F-I, WT(N) and *GAL*-Mia40(N), respectively). They are genetically identical to the strains used so far in this study but were freshly made from a glycerol stock of the wild type to ensure an identical behavior of Rnq1 at the start of the experiment. In both of these naive strains, Rnq1 was only detectable in the pellet fraction (Fig. 5F) and *GAL*-Mia40(N) showed a considerably lower resistance to Q97-GFP than the previous *GAL*-Mia40 strain (Fig. 5G), indicating that the different Rnq1 behavior of these strains strongly contributes to their specific polyQ resistance. However, this *GAL*-Mia40(N) strain could still escape Q97-GFP-mediated growth arrest whereas the wild type strain could not (Fig. 5H). When we cultured these newly made strains in galactose-containing medium (without expression of any polyQ protein), we noticed that over time the proportion of soluble Rnq1 increased, a trend that was considerably more pronounced when Mia40 was overexpressed (Fig. 5I). Thus, the overexpression of Mia40 increases the resistance against polyQ toxicity. This resistance is largely, but not exclusively, exhibited by Mia40-evoqued effects on the formation of Rnq1 seeds (Fig. 5J). This is similar to the situation that was reported before for some cytosolic chaperones, such as Sis1, Sti1 or Hsp104, which likewise reduce polyQ toxicity and likewise reach this at least in part by increasing the solubility of Rnq1 (Higurashi, Hines et al., 2008, Klaips et al., 2020, Meriin et al., 2002, Wolfe et al., 2013).

### Cytosolic aggregates interfere with efficient import of matrix and carrier proteins

How can Mia40 influence the propagation of Rnq1 aggregates in yeast cells? Mia40 is a protein of the mitochondrial IMS and there is no indication for any extramitochondrial fraction of Mia40, not even upon its overexpression. Moreover, overexpression of a cytosolic version of Mia40 (i.e. a variant without its mitochondrial targeting sequence) did not suppress the toxicity of Q97-GFP (Fig. 6A).

**Fig. 6.**
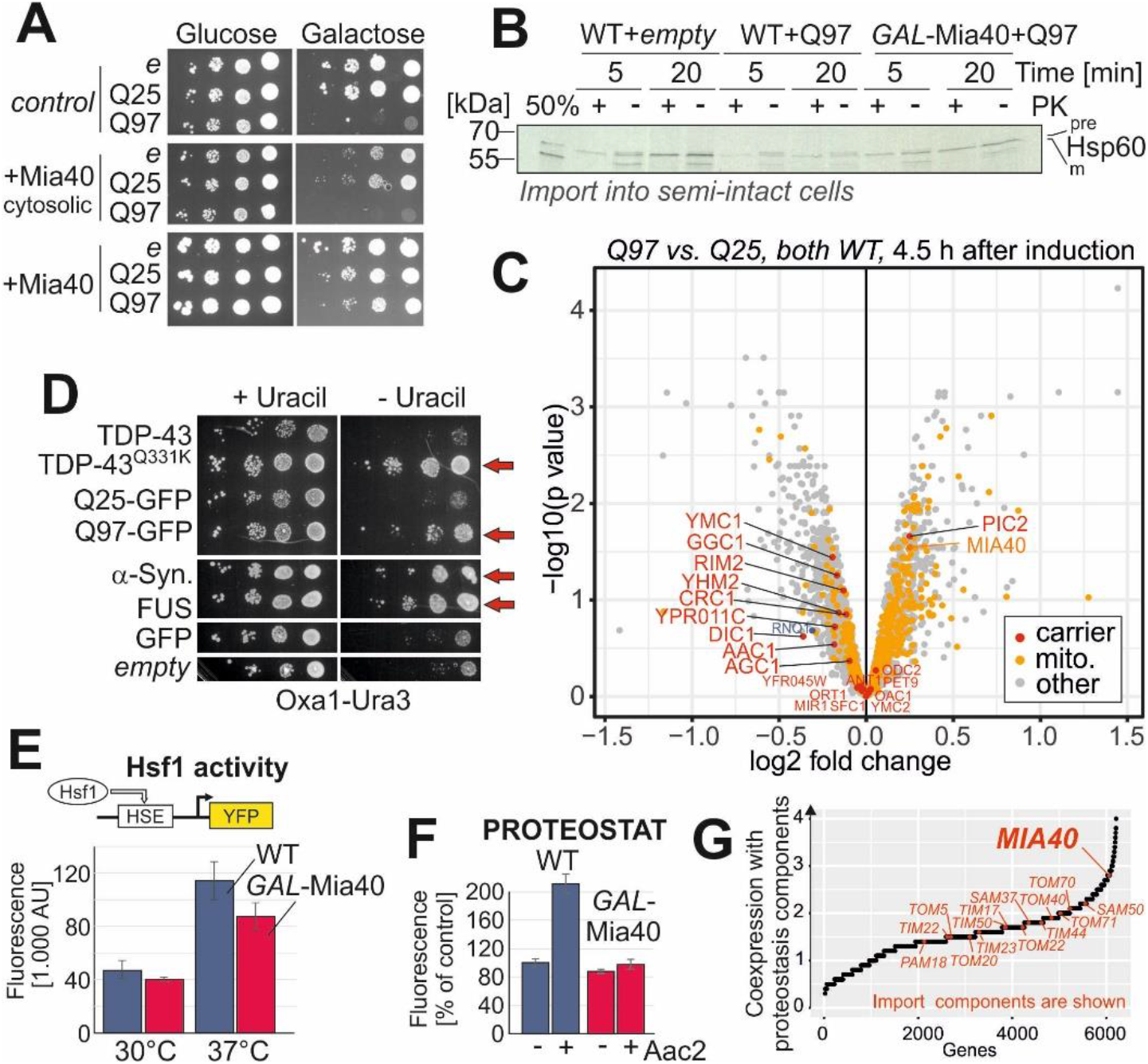
Cytosolic polyQ aggregates interfere with efficient protein import into mitochondria. **A.** Wild type and *GAL*-Mia40 cells, and cells expressing a cytosolic version of Mia40 under *GAL1* control were dropped on the indicated plates. Note, that the mitochondrial version of Mia40 suppressed polyQ toxicity, whereas the cytosolic version did not. **B.** Yeast cells of the indicated strains were converted to semi-intact spheroplasts to which radiolabeled Hsp60 precursor was added. Q97-GFP interfered with the efficient import of Hsp60, particularly in wild type cells. **C.** Q25-GFP and Q97-GFP were expressed in wild type cells for 4.5 h. Cell extracts were subjected to mass spectrometry-based proteomics. The levels of carrier proteins (and Mia40) are visualized in a volcano plot (p-values were derived using a moderated t-test - limma); mito, mitochondrial proteins. **D**. *Δura3* cells expressing the Oxa1-Ura3 reporter for the cytosolic accumulation of mitochondrial precursor proteins (Hansen et al., 2018) were transformed with plasmids expressing the proteins TDP-43, the TDP-43^Q331K^ mutant, Q25-GFP, Q97-GFP, α-synuclein, FUS or GFP. Note that the expression of all aggregation-prone proteins allows cells to grow on uracil-deficient plates indicating the cytosolic accumulation of the Oxa1-Ura3 fusion protein. **E.** Wild type and *GAL*-Mia40 cells were transformed with plasmids that express Yellow Fluorescent Protein (YFP) under the control of a minimal promoter containing the Hsf1-driven heat shock element (HSE). Cells were grown at 30 or 37°C before fluorescence was measured; n=3. **F.** The formation of protein aggregates was detected with a PROTEOSTAT protein aggregation assay. The expression of the most abundant mitochondrial carrier protein, Aac2/Pet9, resulted in a strongly increased fluorescence in wild type cells but not in *GAL*-Mia40 cells. Mean values and standard deviations are shown. **G.** Using the SPELL data collection of transcription data sets, we calculated the co-expression of yeast genes with genes relevant for proteostasis (chaperones, proteasome, and others; see Extended Data for details). Genes for mitochondrial import components are highlighted.

Previous studies showed that cytosolic aggregates can interfere with mitochondrial biogenesis (Cenini et al., 2016, Lehmer, Schludi et al., 2018, Li et al., 2010, Yablonska, Ganesan et al., 2019, Yano et al., 2014). We therefore tested whether the expression of Q97-GFP impaired protein import by the Mia40-mediated pathway. Since only very low levels of Q97-GFP were coisolated with mitochondria, we avoided *in vitro* import experiments with isolated mitochondria but used a previously established protocol employing semi-intact cells (Hansen, Aviram et al., 2018, Laborenz, Hansen et al., 2019). To this end, we removed the cell wall and perforated the plasma membrane by a specific freezing procedure, before radiolabeled precursor of the mitochondrial protein Hsp60 was added. In wild type cells, Hsp60 was efficiently converted to its mature form which was protected against added protease due to its localization within mitochondria. Expression of Q97-GFP considerably impaired the ability of cells to import Hsp60 into mitochondria, an effect that was less pronounced when Mia40 was co-overexpressed (Fig. 6B). In contrast, we did not observe any direct interference of Q97-GFP with the import of Mia40 substrates such as Cox19 (Fig. EV3B).

For a more general and unbiased overview, we quantitatively analyzed the proteomes of wild type cells 4.5 h after induction of either Q25-GFP or Q97-GFP (Fig. 6C). This *in vivo* method indicated a weak, but consistent reduction of carrier proteins, i.e. hydrophobic inner membrane proteins. Taken the relatively short time of polyQ expression (correlating to one or two cell divisions) into account, such a drop in total protein levels could point to a considerable reduction of the biogenesis of these proteins.

We recently developed a sensitive *in vivo* assay to detect the cytosolic accumulation of mitochondrial precursor proteins (Hansen et al., 2018)(Fig. EV3C). Expression of a fusion protein consisting of the nuclear encoded mitochondrial protein Oxa1 and the cytosolic uracil biosynthesis enzyme Ura3 leads to uracil-dependent growth as long as mitochondrial protein import is fully functional, but to uracil independence when import is inefficient (Fig. EV3C). When Oxa1-Ura3 was co-expressed with Q97-GFP, cells became uracil-independent, indicative for the cytosolic accumulation of the Oxa1-Ura3 fusion protein. Similarly, the expression of other aggregation-prone proteins, such as a disease variant of TDP-43, FUS or α-synuclein, also rendered cells uracil-independent, whereas non-aggregating proteins, such as Q25-GFP, TDP-43 or GFP, did not (Fig. 6D). Obviously, cytosolic aggregates of different kind lead to the cytosolic accumulation of the Oxa1-Ura3 reporter, either by interference with its mitochondrial import or with its proteolytic degradation in the cytosol.

Why are elevated Mia40 levels beneficial under these conditions? Previous studies already showed that Mia40 is rate-limiting for the import of its client proteins into mitochondria and that increased Mia40 levels result in increased steady state levels of its substrates (Peleh et al., 2016). Moreover, it was shown that the Mia40-dependent import process directly competes with proteasomal degradation (Kowalski, Bragoszewski et al., 2018, Mohanraj, Wasilewski et al., 2019). It therefore seems conceivable that increased import rates into mitochondria reduce the burden on cytosolic proteostasis. In order to test this, we made use of a reporter system that measures the heat shock response by Hsf1-driven expression of a fluorescent reporter. Growth at 37°C induced the heat shock response to a larger extent in wild type than in *GAL*-Mia40 cells suggesting that this strain indeed is less reactive to heat stress (Fig. 6E).

Presumably owing to their hydrophobic nature, carrier proteins were reported in the past as being a severe burden for cytosolic proteostasis when their mitochondrial import is impaired (Hoshino, Wang et al., 2019, Liu, Wang et al., 2019, Piard, Umanah et al., 2018, Wang & Chen, 2015). We employed an established assay of PROTEOSTAT staining (Pihlasalo, Kirjavainen et al., 2011) to monitor the formation of misfolded proteins in yeast cells. Overexpression of the ATP/ADP carrier Aac2 indeed strongly increased the PROTEOSTAT signal (indicating protein misfolding), but did hardly change the signal in *GAL*-Mia40 cells (Fig. 6G). From this we conclude that high levels of Mia40 can be protective against the toxic effects of mitochondrial precursor proteins. Interestingly, previous studies have shown that cells induce Mia40 when cytosolic precursors accumulate (Boos et al., 2019) and the expression of Mia40 is co-regulated with that of components of the chaperone and proteasome system, in contrast to other mitochondrial import components (Fig. 6G). However, Mia40 overexpression does not generally provide increased stress resistance and can either be protective or sensitizing depending of the stress conditions (Fig. EV3D).

In summary, we propose a model in which the induction of Q97-GFP impairs the biogenesis of a subset of mitochondrial proteins (although details will have to be further studied in the future) and that increased levels of Mia40 might be protective. This effect mechanistically explains why high levels of Mia40 suppress polyQ toxicity.

### Mia40 overexpression also influences Q97-GFP aggregation in a mammalian cell culture model

Is the strong suppression of polyQ toxicity by Mia40 a peculiarity of the yeast model system? To address this question, we first verified that the expression of human MIA40 in human embryonic kidney (HEK293) cells changed the mitochondrial proteome. Western blotting (Fig. 7A) as well as quantitative mass spectrometry (Habich et al., 2019) confirmed that higher levels of MIA40 also increase the steady state levels of some mitochondrial proteins, even though not to the same extent as in yeast (Peleh et al., 2016, Wrobel et al., 2015).

**Figure 7.**
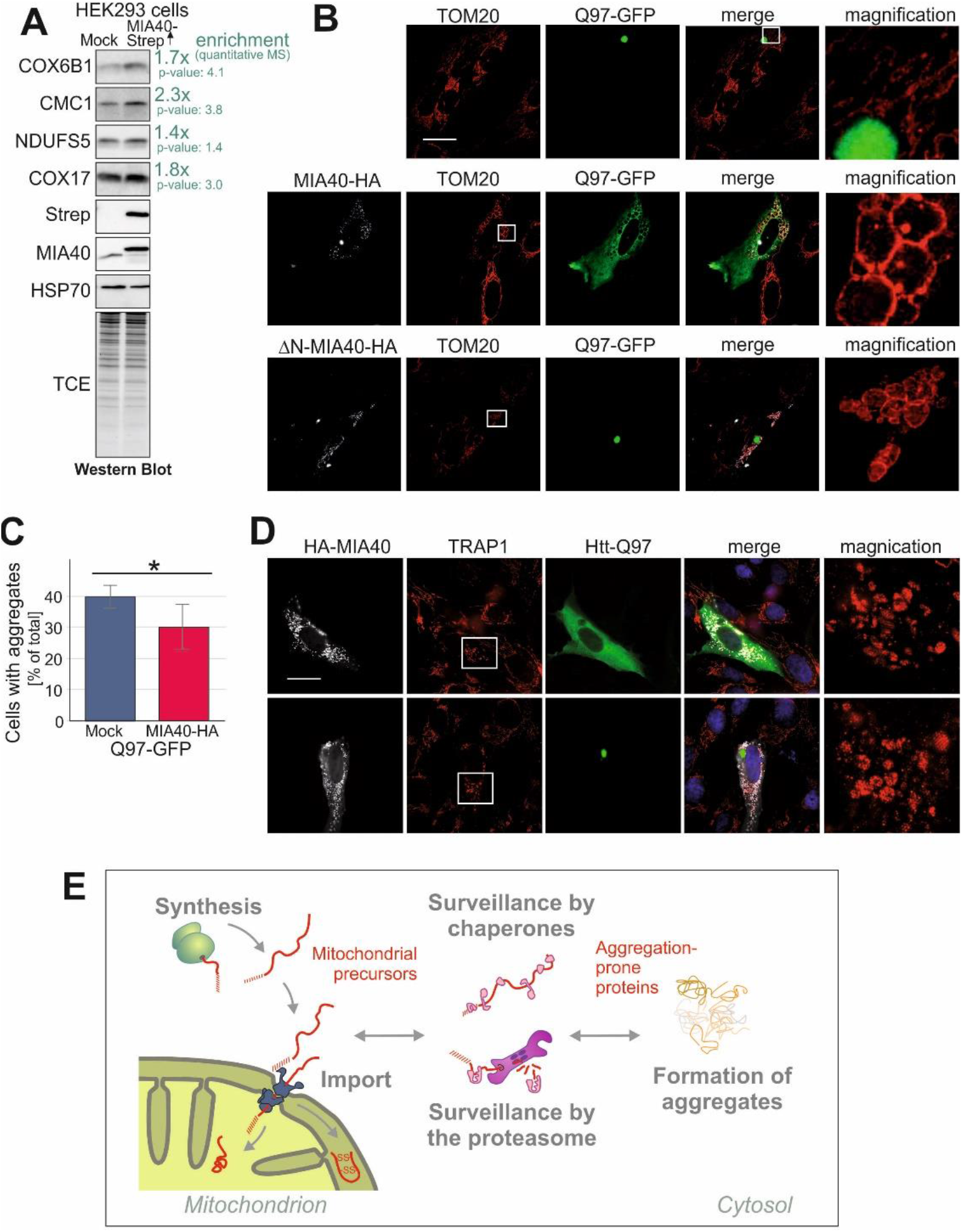
Overexpression of MIA40 reduces the number of human SH-SY5Y cells with polyQ aggregates. **A.** Cell extracts of HEK293 cells expressing a C-terminally Strep-tagged version of MIA40 (MIA40-Strep) or a mock control were analyzed by Western blotting with antibodies against several IMS proteins as well as HSP70 which served as loading control. The enrichment factors shown in green are based on quantitative mass spectrometry data (Habich et al., 2019). **B, D.** SH-SY5Y cells co-expressing Q97-GFP with MIA40 (middle row and D), ΔN-MIA40 (lower row) or a mock-transfected control (upper row) were analyzed by immunocytochemistry and super-resolution microscopy. The magnification shows an enlarged section of the TOM20 staining. TOM20 and TRAP1 serve as markers for the mitochondrial outer membrane and the matrix, respectively. Bars, 20 μm. **C.** The percentage of cells with Q97-GFP aggregates were quantified in control (mock transfected) and MIA40-expressing cells. Data are displayed as mean ± standard deviations and were analyzed by two-tailed Student’s t-test, n = 6.; * P ≤ 0.05. **E.** Model for competition of mitochondrial precursor proteins and aggregation-prone proteins for the same chaperones and components of the proteasome system.

Next, we employed an established cell culture model using human neuroblastoma SH-SY5Y cells that expressed Q97-GFP. This system has been extensively used before as a model for Huntington’s disease (van Well, Bader et al., 2019). In many of these cells, Q97-GFP formed aggregates that often appeared in proximity to mitochondria (Fig. 7B, EV4A). Co-expression of MIA40 in these cells significantly decreased the number of cells with Q97-GFP aggregates (Fig. 7B, C). This suppression of polyQ aggregation was not seen when a variant of MIA40 (ΔN-MIA40) was used which lacked the N-terminal 40 amino acid residues that are important for efficient mitochondrial targeting (Hangen, Feraud et al., 2015). We also noticed that the co-expression of MIA40 and Q97-GFP in SH-SY5Y cells induced changes in the mitochondrial morphology as many mitochondria formed large circular structures (Fig. 7D, EV4B). These structures presumably represent dilated mitochondria as their lumen contained the soluble matrix protein TRAP1. Such alterations were not seen if either MIA40 or Q97-GFP were individually expressed or when MIA40 was co-expressed with Q25-GFP (Fig. EV4C). These morphological changes were reminiscent of the ‘curly’ mitochondria that were induced in yeast cells by co-expression of Mia40 and Q97-GFP (Fig. EV1).

Thus, increased levels of MIA40 can also be protective against the formation of polyQ aggregates in mammalian cells.

## Discussion

Mitochondria can make up a large fraction of the cellular mass (more than 30% in some cell types) and dynamically adapt their volume to metabolic conditions. Owing to their posttranslational mode of import, newly synthesized mitochondrial precursors initially explore the cytosol where they are maintained in a soluble and import-competent state by different chaperones that reside in the cytosol or are associated with the membranes of mitochondria and the ER (Becker, Walter et al., 1996, Deshaies, Koch et al., 1988, Fünfschilling & Rospert, 1999, Hansen et al., 2018, Hoseini, Pandey et al., 2016, Jores, Lawatscheck et al., 2018, Opalinski, Song et al., 2018, Papic, Elbaz-Alon et al., 2013, Terada, Kanazawa et al., 1997). Thereby, mitochondrial protein biogenesis directly depends on the cytosolic chaperone capacity. Presumably as a consequence of their strong tendency to sequester chaperones, the cytosolic accumulation of precursor proteins induces a sudden growth arrest and triggers the increased expression of components of the chaperone and proteasome system (Boos et al., 2019, Boos, Labbadia et al., 2020, Mårtensson, Priesnitz et al., 2019, Shakya, Barbeau et al., 2020, Wang & Chen, 2015, Weidberg & Amon, 2018, Wrobel et al., 2015). Obviously, the post-translational import mode of mitochondrial proteins poses a threat for cellular proteostasis which is met by an adaptive network of cytosolic factors that can deal with unfolded precursors.

Our observations show that the synthesis, targeting, and import of mitochondrial proteins is of direct relevance for the aggregation behavior of cytosolic polyQ proteins. We found that the *mia40-3* mutant is hypersensitive to polyQ proteins even at permissive growth conditions at which no mitochondrial defects are apparent in this strain (Chacinska et al., 2004). Temperature-sensitive Mia40 mutants were instrumental to identify an increased proteasomal capacity as reaction to a reduced mitochondrial import efficiency, a program that is referred to as unfolded protein response activated by mistargeting of proteins (UPRam)(Wrobel et al., 2015). The hypersensitivity of this mutant suggests that the simultaneous presence of accumulated precursors and of polyQ proteins leads to a collapse of proteostasis in these mutants. Apparently, both groups of problematic proteins put a strain on the same components of the chaperone and protein quality control network (Fig. 7E).

It was still surprising to see that the overexpression of Mia40 is protective against cytosolic aggregates. How can this be explained? Two mutually not exclusive mechanisms seem plausible:

1. It was shown in previous studies that Mia40 is rate-limiting for mitochondrial protein import. Increased levels of Mia40 lead to increased protein levels in yeast or human mitochondria (Habich et al., 2019, Peleh et al., 2016, Wrobel et al., 2015). Thus, at normal Mia40 levels, a fraction of newly synthesized mitochondrial proteins is apparently never imported but retained in the cytosol and finally degraded. Recent studies indeed directly demonstrated the competition of the Mia40-mediated import and proteasomal degradation (Kowalski et al., 2018, Krämer, Groh et al., 2020, Mohanraj et al., 2019). Thus, increased Mia40 levels presumably speed up the import of certain proteins and thus might reduce their burden on cytosolic proteostasis.
2. The Mia40 pathway is only one out of several import routes into mitochondria. It is not unlikely that the overexpression of Mia40 might have negative effects on clients of these other import routes. Indeed, we observed that very high levels of Mia40, even higher than those used in this study, are toxic and result in clogger-like effects (Boos et al., 2019, Weidberg & Amon, 2018). Thus, Mia40 overexpression might induce mild perturbations that result in cellular adaptations which give rise to the observed polyQ resistance. Preconditioning to stress conditions is well known from the heat stress response where preexposure to elevated temperature provides resistance to heat. At least in worms, sublethal mitochondrial stress induces a latent heat shock response which is protective against proteostasis collapse (Labbadia et al., 2017, Wu, Senchuk et al., 2018, Zhang, Wu et al., 2018). Such mild Mia40-evoqued stress conditions might also explain the increased elimination of Rnq1 aggregates by counterselection.

Is the modulation of Mia40 levels relevant under physiological conditions? In contrast to other components of the mitochondrial import machinery, the expression of Mia40 is upregulated upon stress conditions induced by mitochondrial defects (Boos et al., 2019, Zöller, Laborenz et al., 2020) but also by problems outside mitochondria (Metzger & Michaelis, 2009). Transcription of *MIA40* is under control of Rpn4, a transcription factor that adjusts proteasome levels (Xie & Varshavsky, 2001). Most substrates of Mia40 are small hydrophilic proteins of the IMS that are unlikely to be harmful for the cytosol. However, major clients of the Mia40 pathway are the small Tim proteins, essential components for the import of inner membrane carriers (Koehler, Jarosch et al., 1998, Luciano, Vial et al., 2001). Carriers are very abundant hydrophobic proteins and it is well documented that misfolded or non-imported carriers pose a major threat to cytosolic proteostasis (Hoshino et al., 2019, Liu et al., 2019, Vartiainen, Chen et al., 2014, Wang & Chen, 2015). Consistent with this idea, a previous genome-wide overexpression screen for suppressors of polyQ toxicity identified a large number of components of the carrier import pathway, including Erv1, Tim9, Tim10, Tim12, and Tim22 (Mason et al., 2013). It will be very exciting to unravel the specific impact of distinct precursor proteins in the context of cytosolic proteostasis.

Our study documents that modulation of the capacity of the mitochondrial import machinery is of direct relevance for cytosolic proteostasis. Upregulation of mitochondrial import components, in particular of Mia40, might provide a strategy to stabilize aggregation-prone proteins in human cells. The well-known decline of cytosolic proteostasis in aging cells, and in particular in neurons of patients with Huntingon’s disease, along with slower import rates into mitochondria would increase the abundance of cytosolic precursors and the occupancy of precursor-binding chaperones. The upregulation of Mia40 levels or ways to increase the functionality of the mitochondrial disulfide relay may provide a strategy to better stabilize cytosolic proteostasis in such cells and will be exciting to be explored in the future.

## Material and Methods

### Yeast strains and plasmids

All yeast strains used in this study are based on the wild type strains BY4742 (MATα *his3 leu2 lys2 ura3*) and YPH499 (MATa *ura3 lys2 ade2 trp1 his3 leu2*). They are listed in Table EV1. The *mia40*-3 mutant (Chacinska et al., 2004) and the *GAL*-Mia40 (Terziyska, Lutz et al., 2005) strain were described previously. For the generation of *GAL*-Mia40(N), the sequence of the *HIS3* gene and the *GAL10* promoter were amplified from the plasmid pTL26 and inserted upstream of *MIA40*. To delete *RNQ1* in YPH499 and *GAL*-Mia40, the kanMX4 cassette for G418 resistance was amplified from pFA6a-kanMX4 and genomically integrated by homologous recombination.

For the Hsf1 activity reporter, the pNH605-4xHSEpr-YFP reporter plasmid was used as described before (Zheng, Krakowiak et al., 2016). To generate an *AAC2* overexpression plasmid, the coding region of *AAC2* was amplified from genomic DNA and ligated into a pYX233 empty vector plasmid between the *GAL1* promoter and an HA tag by using the restriction sites *Eco*RI and *Xma*I. To generate pET19b-Rnq1, the *RNQ1* ORF was amplified from isolated YPH499 gDNA using primers for the introduction of *Nde*I and *Bam*HI restriction sites. After restriction digest of the PCR product and the pET19b vector, ligation was performed to generate the pET19b-Rnq1 plasmid for the bacterial expression of N-terminally His-tagged Rnq1.

To generate Entry Clones and Expression Clones, the Gateway technology was applied according to the instructions of the manufacturer (Invitrogen). BP reactions and LR reactions were used basically as described in the Gateway technology user guide. To generate Entry Clones, the ORF of interest was amplified by PCR using YPH499 genomic DNA as template and primers for the introduction of *att*B-sites. The resulting *att*B-PCR product was mixed with the pDONR221 Donor Vector in a BP reaction. To generate Expression Clones, the Entry Clone carrying the ORF of interest was mixed with the Destination Vector containing the desired properties for expression in yeast in an LR reaction.

To generate a non-fluorescent version of Q97-EGFP, the glycine at position 67 of the EGFP amino acid sequence was changed to alanine in the respective Entry Clone by site-directed mutagenesis. To this end, primers to introduce a G to C mutation were designed to be 28 nucleotides long and to carry the desired mutation in the middle of the primer. The Entry Clone was then subjected to error-prone PCR using the indicated primers and the PfuUltra HF DNA polymerase. The template DNA was removed after the reaction by digestion with *Dpn*I. The same cloning strategy was used to generate a non-fluorescent version of Q25-EGFP. After sequencing of the Entry Clones, an LR reaction was performed to produce the pAG424GAL-Q25-EGFPnF and pAG424GAL-Q97-EGFPnF plasmids.

Strains were grown in yeast complete medium (1% yeast extract, 2% peptone), containing 2% of the carbon sources galactose or glucose as indicated. The temperature-sensitive strain (*mia40-3*) was grown at 25°C before switching it to 30°C. Strains containing plasmids were grown at 30°C in minimal synthetic medium containing 0.67% yeast nitrogen base and 2% lactate as carbon source. To induce the expression from the *GAL1* promoter, cultures were supplemented with 2% galactose.

### Human cell lines and plasmids

For transfection, 0.15 to 0.3 μg of DNA per 24-well (150 000 cells plated) was mixed with 50 μl Opti-MEM and 4 μl polybrene in tube A. Meanwhile, 1 μl lipofectamine was added to tube B which also contained 50 μl Opti-MEM. Both tubes were incubated for 15 minutes. The contents of both tubes were and further incubated for 15 min. Cells were washed with PBS (−/−) and 500 μl of Opti-MEM was added to each well. Following the 15 min incubation, 100 μl of the mix was added dropwise to each well.

### Data Availability

The mass spectrometry proteomics data have been deposited to the ProteomeXchange Consortium via the PRIDE (Perez-Riverol, Csordas et al., 2019) partner repository.

### Miscellaneous

The following methods were performed according to published methods: Isolation of mitochondria and oxygen consumption measurements (Saladi, Boos et al., 2020); analysis of mRNA levels by qRT-PCR (Zöller et al., 2020); preparation of semi-intact cells and their use for *in vitro* import experiments (Laborenz et al., 2019); pulse chase assay and alkylation for *in vivo* analysis of Mia40-mediated import (Peleh et al., 2016); the analysis of mitochondrial protein import using the Oxa1-Ura3 reporter assay (Hansen et al., 2018); growth and analysis of MIA40-expressing HEK293T cells (Fischer, Horn et al., 2013, Murschall, Gerhards et al., 2020).

For further information on materials and methods, please consult the Appendix that is provided in the Expanded View.

## Supporting information

Supplemental Table 1 - Strains

Supplemental Table 2 - Proteomics

Supplemental Table 3 - Antibodies

## Acknowledgements

We thank Sabine Knaus, Andrea Trinkaus and Laura Buchholz for technical assistance, Agnieszka Chacinska for the *mia40-3* mutant stain, Kai Hell for the *GAL*-Mia40 strain, Flaviano Giorgini for the *GALL*-Q103GFP plasmid, Katja G. Hansen for the Oxa1-Ura3 plasmid and Nils Wiedemann and Nikolaus Pfanner for antibodies. This study was funded by grants from the Deutsche Forschungsgemeinschaft (DIP MitoBalance, SPP1710, IRTG1830, HE2803/10-1 to JMH, CRC1218/B02 to JR, WE2174/7-1 to BW, and WI/2111-6 and WI/2111/8 to KFW), the Landesschwerpunkt BioComp (to JMH), the Joachim Herz Stiftung (to FB) and the Elitenetzwerk Bayern (Biological Physics program, to JS). SR-SIM microscopy was funded by the Deutsche Forschungsgemeinschaft and the State Government of North Rhine-Westphalia (INST 213/840-1 FUGG).

## Author contributions

A.M.S. and K.K. designed, cloned and verified the constructs and strains; J.M.H. conceived the project; A.M.S. and K.K. characterized the role of Mia40 in the context of polyQ-induced stress in yeast; S.M. and G.C. analyzed the polyQ aggregation by microscopy using microfluidics chambers and live cell imaging; N.K. and S.A. programmed the automated detection of aggregation patterns using machine learning; J.S., R.J.B. and B.W. analyzed the influence of polyQ aggregates and Mia40 on mitochondrial morphology in yeast; A.S.V., J.R. and K.F.W. planned and carried out the experiments in human cells to analyze the relevance of MIA40 in the context of polyQ aggregation; P.H., F.S.,F.B. and A.M.S. analyzed the cellular proteome by mass spectrometry; all authors analyzed the data; J.M.H. wrote the manuscript with assistance from all other authors.

## Conflict of interest

The authors declare that they have no conflict of interest.

## EXTENDED VIEW

**Figure EV1.**
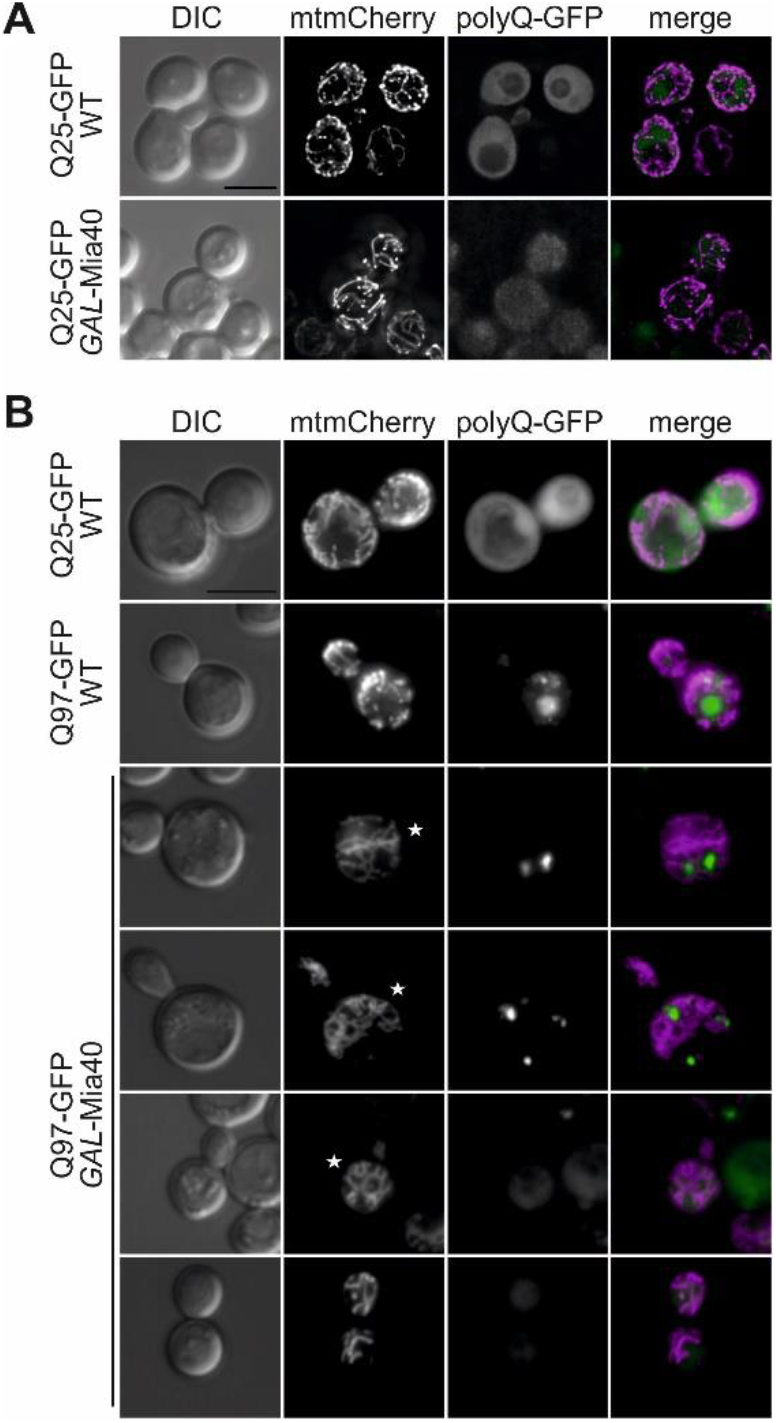
Co-expression of Q97-GFP and *GAL*-Mia40 results in minor changes of mitochondrial morphology. **A.** Cells were grown to mid-log phase in glucose-containing medium, shifted to galactose-containing medium for 4 h, and analyzed by 3D fluorescence microscopy. Fluorescence micrographs are z stacks subjected to deconvolution. DIC, differential interference microscopy. Bar, 5 μm. **B.** Cells were grown to mid-log phase in medium containing glycerol (3%) and ethanol (2%) as carbon sources, shifted to galactose-containing medium for 4 h, and analyzed by 3D fluorescence microscopy. Fluorescence micrographs are maximum intensity projections of z stacks. Asterisks indicate representative cells exhibiting interconnected, ‘curly’ mitochondria. Bar, 5 μm.

**Figure EV2.**
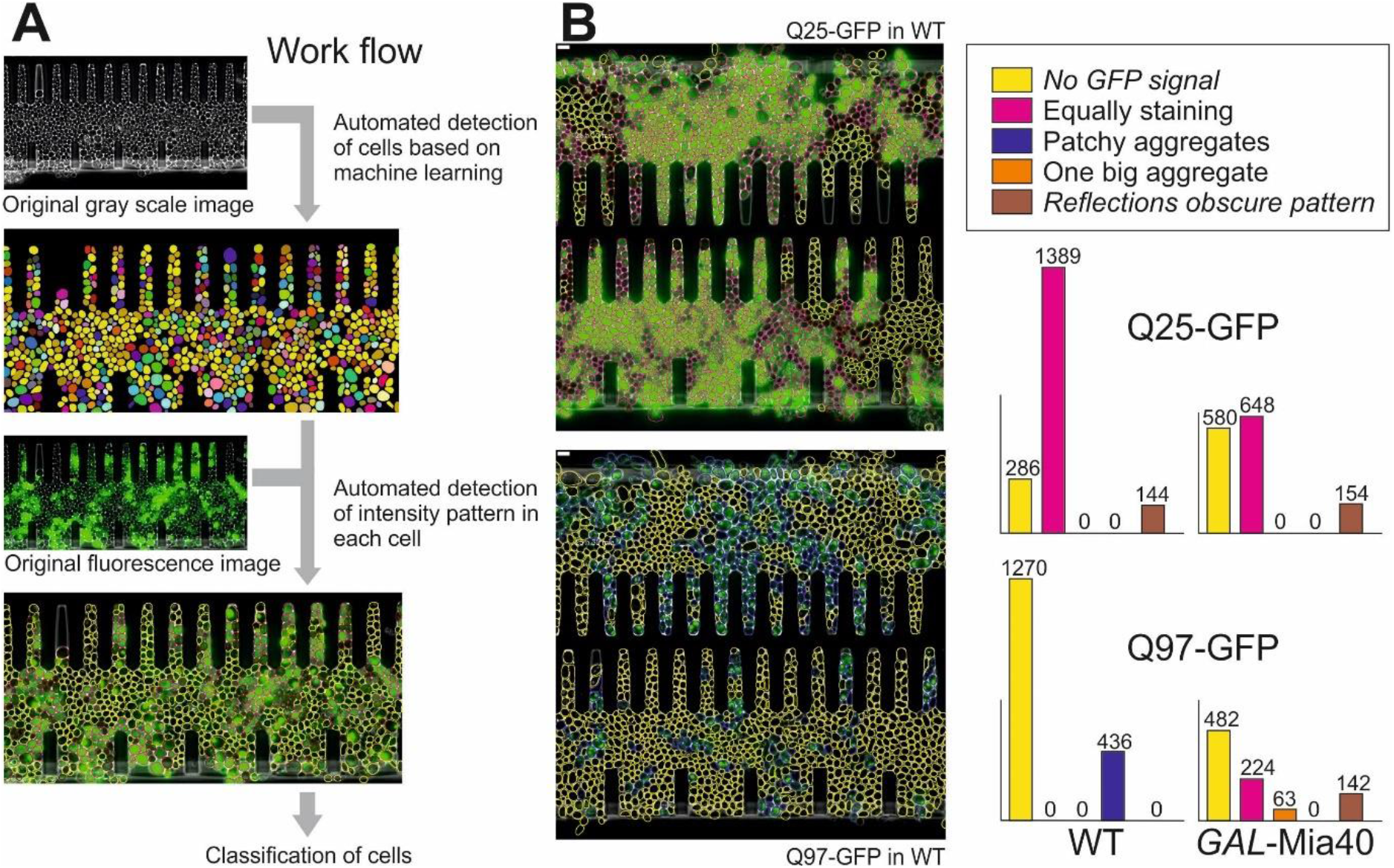
The different patterns of aggregate formation can be quantified by an automated method trained with a machine learning approach. **A.** An automated detection algorithm was established that was trained by use of 154 original gray scale images of growing yeast cells for which each cell had been outlined by hand by visual inspection. Machine learning optimized cell detection before the patterns of fluorescence were again detected manually in a training set from which the algorithm learned to distinguish five different classes of patterns. **B.** Examples of these automatically detected classes of cells are shown in which the five categories are indicated by different colors. The number of cells in each category are shown here for different yeast strains as indicated.

**Fig. EV3.**
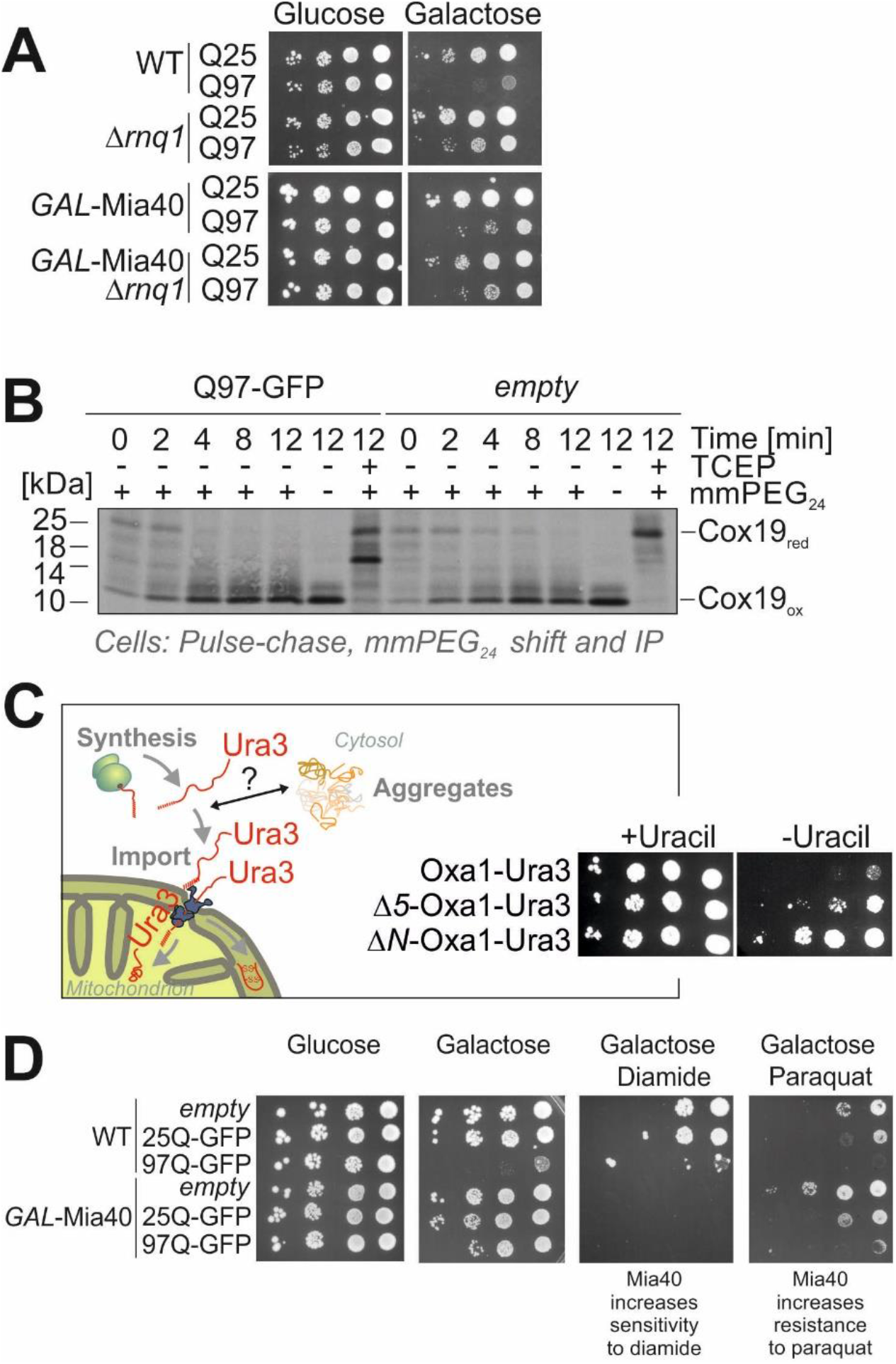
Increased levels of Mia40 can have positive or negative effects, depending on the specific conditions. **A.** Drop dilution experiment of the indicated strains. **B.** Wild type cells containing the Q97-GFP-expression or empty plasmids were radiolabeled for 3 min with ^35^S-methionine before labeling was quenched by addition of excess amounts of non-radioactive methionine. At the time points indicated, proteins were precipitated with trichloroacetic acid before the alkylating agent methyl-polyethylene glycol (24)-maleimide (mmPEG24) was added which shifts the size by about 2 kDa per free reduced thiol group. For control, tris(2-carboxyethyl) phosphine (TCEP) was added to reduce all disulfide bonds. **C.** Wild type cells carrying the *Δura3* marker allele were transformed with plasmids for the expression of Oxa1-Ura3, Δ5-Oxa1-Ura3 (lacking the N-terminal five residues) or ΔN-Oxa1-Ura3 (lacking the entire 42 residues of the mitochondrial targeting sequence of Oxa1). Serial dilutions were dropped on plates containing or lacking uracil. **D.** The indicated strains were grown to mid-log phase in glucose medium before serial dilutions were dropped on the indicated plates and incubated at 30°C.

**Figure EV4.**
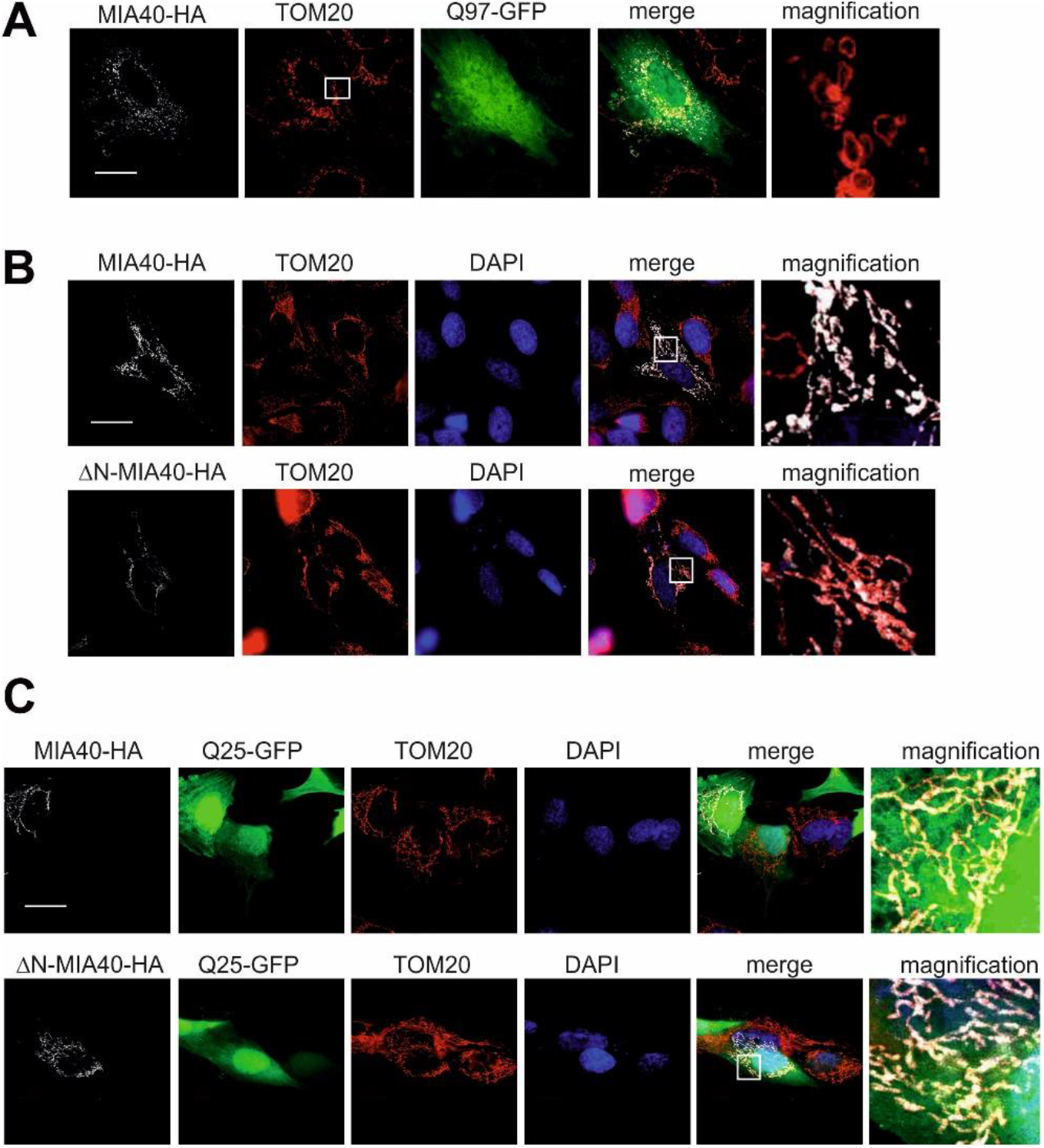
The expression of MIA40, ΔN-MIA40 or Q25-GFP do not affect mitochondrial morphology. **A-C.** SH-SY5Y cells were analyzed by immunocytochemistry as indicated. TOM20 was used as mitochondrial marker. Whereas MIA40 colocalizes with TOM20, ΔN-MIA40 does not or is associated with the mitochondrial surface. Bars, 20 μm.

## Appendix Expanded View – Appendix

## Additional information to Material and Methods

## Culture conditions of human cells

SH-SY5Y cells were kept in a humidified incubator at 37°C, 5% CO2. Passage of the cells was performed as follows: cells were washed with PBS (−/−), then 3 ml of trypsin were added for a 175 cm2 flask and incubated for 2 minutes in the incubator. Trypsinized cells were resuspended in 7 ml medium (DMEM F12 + 15 % fetal calf serum + 1 % penicillin/streptomycin + 1 % non-essential amino acids (NEAA); obtained from Gibco, Invitrogen) and centrifuged. The cell pellet was again resuspended in 10 ml medium and divided in flasks in a proportion of 1/10.

## Viability assay

Yeast cells were grown to mid log phase in lactate medium before the induction of Q97-GFP was induced by the addition of 2% galactose. Samples were taken after different times and diluted to OD (600 nm) of 0.0002 in 300 μl of sterile water. 80 μl of the cell suspension was plated in technical triplicates on glucose-containing plates to repress the expression of Q97-GFP. The plates were incubated for 5 days to assess the number of colonies which survived the expression of Q97-GFP.

## Subcellular fractionation analysis of Rnq1

To determine the aggregation state of Rnq1, a previously established protocol was followed (Sideri, Koloteva-Levine et al., 2011). Cells were grown until log phase. 5.0 OD (600 nm) were harvested and washed once with distilled water. The cell pellet was resuspended in 200 μl of lysis buffer (74 mM Tris-HCl pH 8.0, 150 mM KCl, 50 mM EDTA, 1 mM DTT, 0.2% SDS, 1% Triton X-100, 1 mM PMSF, 1x Complete mini protease inhibitor mixture (Roche)) and transferred to a new tube containing 200 μl of glass beads for mechanical lysis using a Fastprep bead beater. To remove cell debris, the lysates were centrifuged for 3 min at 4°C and 3,000 rpm. 50 μl of the supernatant were added to 50 μl of reducing sample buffer as the total. Another 50 μl were transferred to a new tube and centrifuged for 1 h at 85,000 x g, 4°C, to generate supernatant and pellet fractions. The supernatant was transferred to a new tube and the pellet was resuspended in 50 μl of lysis buffer. 50 μl of reducing sample buffer were added to both, the supernatant and the pellet fraction. All samples were incubated at 96°C for 4 min and subsequently subjected to SDS-PAGE and Western Blot using an anti-Rnq1 antibody for the detection of Rnq1 in either the pellet or the supernatant fraction.

## Sample preparation, mass-spectrometry and proteomics data analysis

To compare the soluble proteomes of wild type and *GAL*-Mia40 yeast strains which both have been expressing Q97-GFP for 4.5 h, as well as the changes in the proteomes of WT strains expressing either Q97-GFP or Q25-GFP, the expression of the polyQ constructs was induced for 0 h and 4.5 h. 50 ml of mid-log phase cell suspension were harvested by centrifugation at RT, 4,000 x g for 5 min and washed twice with PBS before freezing the pellet in liquid nitrogen and storing it at −80°C. The cells were resuspended in PBS with a volume in ml equal to 0.4 x OD_600_ (e.g., if OD_600_=0.5, add 200 μl; calculation: 0.5 x 0.4 = 0.2 ml). 40 μl of the cell suspension were transferred to a new tube to which 60 μl of cold 1.7x lysis buffer (0.8% NP-40 in PBS, 1 mM MgCl_2_, 0.25 U/μl benzonase, complete Protease Inhibitor Cocktail, Phosphatase Inhibitor, PBS) were added. Cell lysates were prepared using a FastPrep-24 5G homogenizer (MP Biomedicals, Heidelberg, Germany) with 3 cycles of 30 s, speed 8.0 m/s, 120 s breaks, glass beads. To remove cell debris, the lysates were centrifuged at 4°C for 10 min at 16,100 x g. 40 μl of the supernatant were transferred onto a pre-wet 0.45 μm filter plate on a 96-well plate and centrifuged for 5 min at 500 x g, 4°C to collect the lysate free of Q97-GFP aggregates. The sample plate was sealed with a plate seal plastic foil, dipped in liquid nitrogen and frozen at −80°C until use.

For LC-MS/MS analysis of lysates, protein concentration of lysates were determined by BCA protein determination. 10 μg of each lysate were subjected to an in-solution tryptic digest using a modified version of the Single-Pot Solid-Phase-enhanced Sample Preparation (SP3) protocol (Hughes, Foehr et al., 2014, Moggridge, Sorensen et al., 2018). Lysates were added to Sera-Mag Beads (Thermo Scientific) in 10 μl 15% formic acid and 30 μl of ethanol. Binding of proteins was achieved by shaking for 15 min at room temperature. SDS was removed by 4 subsequent washes with 200 μl of 70% ethanol. Proteins were digested overnight at room temperature with 0.4 μg of sequencing grade modified trypsin (Promega) in 40 μl Hepes/NaOH, pH 8.4 in the presence of 1.25 mM TCEP and 5 mM chloroacetamide (Sigma-Aldrich). Beads were separated, washed with 10 μl of an aqueous solution of 2% DMSO and the combined eluates were dried down. Peptides were reconstituted in 10 μl of H_2_O and reacted for 1 h at room temperature with 80 μg of TMT10plex (Thermo Scientific) (Werner, Sweetman et al., 2014) label reagent dissolved in 4 μl of acetonitrile. Excess TMT reagent was quenched by the addition of 4 μl of an aqueous 5% hydroxylamine solution. Peptides were reconstituted in 0.1 % formic acid, mixed to achieve a 1:1 ratio across all TMT-channels and purified by a reverse phase clean-up step (OASIS HLB 96-well μElution Plate, Waters). Peptides were subjected to an off-line fractionation under high pH conditions (Hughes et al., 2014). The resulting 12 fractions were then analyzed by LC-MS/MS on an Orbitrap Fusion Lumos mass spectrometer (Thermo Scientific) as previously described (Sridharan, Kurzawa et al., 2019). To this end, peptides were separated using an Ultimate 3000 nano RSLC system (Dionex) equipped with a trapping cartridge (Precolumn C18 PepMap100, 5 mm, 300 μm i.d., 5 μm, 100 Å) and an analytical column (Acclaim PepMap 100. 75 × 50 cm C18, 3 mm, 100 Å) connected to a nanospray-Flex ion source. The peptides were loaded onto the trap column at 30 μl per min using solvent A (0.1% formic acid) and eluted using a gradient from 2 to 40% Solvent B (0.1% formic acid in acetonitrile) over 2 h at 0.3 μl per min (all solvents were of LC-MS grade). The Orbitrap Fusion Lumos was operated in positive ion mode with a spray voltage of 2.4 kV and capillary temperature of 275 °C. Full scan MS spectra with a mass range of 375–1500 m/z were acquired in profile mode using a resolution of 120,000 (maximum fill time of 50 ms or a maximum of 4e5 ions (AGC) and a RF lens setting of 30%. Fragmentation was triggered for 3 s cycle time for peptide like features with charge states of 2–7 on the MS scan (data-dependent acquisition). Precursors were isolated using the quadrupole with a window of 0.7 m/z and fragmented with a normalized collision energy of 38. Fragment mass spectra were acquired in profile mode and a resolution of 30,000 in profile mode. Maximum fill time was set to 64 ms or an AGC target of 1e5 ions). The dynamic exclusion was set to 45 s.

Acquired data were analyzed using IsobarQuant (Franken, Mathieson et al., 2015) and Mascot V2.4 (Matrix Science) using a reverse UniProt FASTA Saccharomyces cerevisiae database (UP000002311) including common contaminants and the two protein sequences of Q25-GFP and Q97-GFP.

The following modifications were taken into account: Carbamidomethyl (C, fixed), TMT10plex (K, fixed), Acetyl (N-term, variable), Oxidation (M, variable) and TMT10plex (N-term, variable). The mass error tolerance for full scan MS spectra was set to 10 ppm and for MS/MS spectra to 0.02 Da. A maximum of 2 missed cleavages were allowed. A minimum of 2 unique peptides with a peptide length of at least seven amino acids and a false discovery rate below 0.01 were required on the peptide and protein level (Savitski, Wilhelm et al., 2015).

The raw output files of IsobarQuant (protein.txt – files) were processed using the R programming language (ISBN 3-900051-07-0). Only proteins that were quantified with at least two unique peptides were considered for the analysis. Moreover, only proteins which were identified in both mass spec runs were kept. 2915 proteins passed these quality control filters. Raw TMT reporter ion signals (signal_sum columns) were first cleaned for batch effects using using the ‘removeBatchEffect’ function of the limma package (Ritchie, Phipson et al., 2015) and further normalized using vsn (variance stabilization normalization) (Huber, von Heydebreck et al., 2002). Missing values were imputed with knn method using the Msnbase package (Gatto & Lilley, 2012). Proteins were tested for differential expression using the limma package. T-value outputs of the limma package were used as an input for the fdrtool function of the fdrtool package (Strimmer, 2008) in order to estimate p-values and false discovery rates (qvalues were used). Proteins were classified as ‘hit’ with an fdr smaller 5% and a fold-change of at least 50% and classified as ‘candidate’ with an fdr smaller 20% and a fold-change of at least 40% using either the fdr estimated by limma directly or by the fdrtool package, depending on which method led to a faster approach to zero of the fdr with increasing effect size (t value).

## Hsf1 activity reporter assay

The cells were grown to log phase (OD600 value of 0.4-0.8) in YPGal at 30°C. Part of every culture was incubated for 4 h before the measurement was started at 37°C. Equal amounts of cells (5.0 OD (600 nm)) were collected by centrifugation (4,000 x g, 5 min, room temperature) and resuspended in 666.6 μl MES-Tris pH 6.8. 200 μl of this cell suspension (1.5 OD (600 nm)) were transferred to a flat-bottomed black 96-well imaging plate (BD Falcon) in technical replicates. Cells were sedimented by gentle spinning (30 x g, 5 min, room temperature) and fluorescence (excitation 497 nm, emission 540 nm) was measured using a ClarioStar Fluorescence plate reader (BMG-Labtech).

## PROTEOSTAT staining

To assess the occurrence of protein aggregates in cells, the cells were grown in lactate selective media to log phase, then supplemented with 2% galactose to induce the overexpression of Aac2 for 4.5 h. Then, 5.0 OD (600 nm) were harvested and fixed by the addition of formaldehyde to a final concentration of 4% in 1 ml of sterile water for 15 min. After washing two times with PBS, the cells were resuspended in 1 ml PROTEOSTAT (Enzo Life Sciences GmbH) buffer and incubated at room temperature for 30 min. After centrifugation, the cell pellet was resuspended in 250 μl PROTEOSTAT reagent (1:2000 diluted in PROTEOSTAT buffer) and incubated for 30 min at 4°C. The cells were washed with 500 μl of buffer, then resuspended in 500 μl of buffer. 100 μl of this cell suspension was transferred to a flat-bottomed black 96-well imaging plates in technical replicates. Cells were sedimented by gentle spinning (30 x g, 5 min, room temperature) and fluorescence (excitation, 550 nm; emission, 600 nm) was measured using a ClarioStar Fluorescence plate reader (BMG-Labtech).

## Coexpression analysis of mitochondrial import components with the cytosolic proteostasis system

To analyze the coexpression of mitochondrial import components with components of the proteostasis network, the Serial Pattern of Expression Levels Locator (SPELL) database (Hibbs, Hess et al., 2007) was queried for the set of major cytosolic stress-reactive proteostasis factors (*SSA1, SSA2, SSA3, SSA4, HSP82, HSC82, HSP42, HSP104, HSP26, YDJ1, XDJ1, CCT2, CCT3, PRE1, PRE2, PUP1, RPN1, RPN2, RPT1, RPT2, CDC48)* with the online interface provided by the *Saccharomyces* genome database (SGD, https://spell.yeastgenome.org). All genes in the yeast genome were ranked according to their coexpression score across all transcriptomics datasets deposited to SGD and mitochondrial import components were highlighted.

## Protein purification and antibody production

To generate an antibody recognizing Rnq1, the *RNQ1* ORF was cloned into the expression plasmid pET19b using NdeI and BamHI restriction sites for the bacterial expression of N-terminal His_6_-tagged Rnq1. This recombinant protein was expressed in *E. coli* Rosetta cells at 37°C. Once growth reached an OD (600nm) value of 0.45, the cells were subjected to induction by 1 mM IPTG for 2 h. Cells were harvested by centrifugation and lysed in 400 μl buffer L1 (25% sucrose 50mM Tris/HCl pH 8.0) for 10 minutes. Then, 17.4 ml L1 were added and supplemented with 25 mM EDTA, 1% Triton X-100, 1 mM PMSF, 10 mM DTT. The lysate was frozen overnight at −70°C. Subsequently, the lysates were sonicated three times for 10 seconds and centrifuged at 47,000 x g for 30 minutes. After resuspending the pellet in 10 ml L2 buffer (20 mM Tris/HCl pH 7.4, 1 mM EDTA, 1 mM PMSF, 1% Triton X-100, 50 mM DTT), another sonication step followed by centrifugation was performed. The resulting pellet was resuspended in 10 ml buffer L3 (L2 + 0.1% Triton X-100), sonified and centrifuged. Then, the pellet was resuspended in 10 ml buffer L4 (L2 without Triton X-100), sonicated and centrifuged, and afterwards resuspended in 1 ml buffer L5 (7 M urea, 50 mM Tris/HCl pH 7.4, 50 mM DTT). After a final sonication step, the lysate was frozen at −20°C until it was used for the immunization of rabbits.

## Antibodies

Antibodies used in this study are listed in Table EV3. Antibodies for the use in *S. cerevisiae* were raised in rabbits using recombinant purified proteins. The secondary antibody was ordered from BioRad (Goat Anti-Rabbit IgG (H+L)-HRP Conjugate). The horseradish-peroxidase coupled HA antibody for western blotting was obtained from Roche (Anti-HA-Peroxidase, High Affinity (3F10)). Antibodies were diluted in 5% nonfat dry milk-TBS.

## Immunocytochemistry

48 hours after transfection, cells were washed twice in PBS (−/−) and fixed in 4% PFA for 10 minutes at room temperature. Then, samples were incubated with blocking solution (5% goat serum in 0.2% Triton X-100) for two hours at room temperature. Afterwards, the primary antibody was prepared in blocking solution at the indicated concentration and incubated overnight at 4°C in a humidified environment. The next day, coverslips were washed twice in PBS (−/−) and incubated with secondary antibody in PBS (−/−) for 2 h at room temperature. Then, coverslips were washed twice in PBS (−/−) for 10 minutes, twice in 0.2% Tween in PBS and twice in PBS (−/−) for 10 minutes. Finally, cells were stained with DAPI and mounted on coverslips with Fluoromount medium (Thermofisher).

## Coating of coverslips

Poly-L-lysine (25 μg/ml in PBS −/−) was applied to autoclaved coverslips and incubated for 1 h. Then, coverslips were washed twice with PBS and laminin solution (4 ng/ml in PBS +/+) was added and incubated overnight in the incubator. The next day, the glass coverslips were washed twice in PBS and cells were seeded.

## Microscopic analysis

To analyse the formation of aggregates in yeast, mid log phase cultures were shifted to media containing galactose to induce the expression of Q97-GFP and incubated at 30°C for different times. After centrifugation of 1.2 OD (600 nm) of cells (1 min at 16,100 x g at room temperature), the cells were washed and resuspended in 250 μl of sterile water. Subsequently, the cell suspension was fixed on a microscope slide for fluorescence imaging using the 100x oil objective of a Nikon Eclipse E600 Microscope. Microscopy images were processed in Fiji.

For fluorescence microscopy and quantification of different aggregation phenotypes at different time points, yeast cells were incubated 1.5 - 24 h under inducing conditions. 1 ml of cell culture was pelleted, washed and resuspended in 100 μl of sterile water. To assess the nature of the aggregation pattern in the different strains, 100 cells of each sample were counted and evaluated regarding the aggregation pattern they exhibited.

To assess the cellular localization of Q97-GFP and mitochondrial morphology in the presence of Q97-GFP aggregates in yeast, cells were incubated under non-expressing conditions at 30°C for 16 h. For induction of Q97-GFP expression, cells were then shifted to galactose medium, diluted two-fold and incubated at 30°C for 4 h. Living cells were analysed using a Leica DMi8 fluorescence microscope (Leica Microsystems GmbH, Wetzlar, Germany) with HC PL APO 100x/1.40 OIL objektive, a Lumencor SPECTRA X light source and fluorescence filter sets (FITC ex. 460-500 nm; em. 512-542 nm and TXR ex. 540-580 nm; em. 592-668 nm). The microscope was equipped with a sCMOS Leica-DFC9000GT-VSC07400 camera. Voxel size of the shown images is 0.066 μm. For microscope settings, image generation and processing (cropping, maximum intensity projection) the Leica LAS X software (version 3.4.2.18368, Leica Microsystems GmbH, Wetzlar, Germany) was used. Deconvolution of z stacks was carried out by Huygens Deconvolution Software (Scientific Volume Imaging, Hilversum, The Netherlands). Final image processing, including adjustment of brightness, contrast and background reduction, and the overlay of different channels was done using Adobe Photoshop CS5 (Adobe Systems).

To perform microfluidics, microfluidic chips were made as previously described (Goulev, Morlot et al., 2017, Morlot et al., 2019). The micro channels were cast by curing PDMS (Sylgard 184, 10:1 mixing ratio) and then covalently bound to a 24×50 mm coverslip using plasma surface activation (Diener, Germany). The assembled chip was then baked for 1 h at 60°C and then perfused with media using Tygon tubing and a peristaltic pump (Ismatec, Switzerland) at a 10 μl/min flow rate. After 2 h of PDMS rehydration, yeast cells were loaded into the chip with a 1 ml syringe. Cells were grown on chip with constant perfusion of glucose selective medium (SD-Leu-Ura) until most cavities were filled with cells (typically overnight). Then induction was started by changing the medium to SC+2%Galactose-Leu-Ura. At the same time, time-lapse acquisition was initiated. For the glucose recovery experiments, medium was switched back to SD-Leu-Ura after 22 h acquisition of galactose induction. For all other experiments, the same medium (SC+2%Galactose-Leu-Ura) was perfused until the end of the time-lapse. To this end, cells were imaged using an inverted Nikon Ti-E microscope. Fluorescence illumination was achieved using LED light (Lumencor) and emitted light was collected using a 60x N.A. 1.4 objective and a CMOS camera Hamamatsu Orca Flash 4.0. An automated stage was used to follow up to 34 different fields of view in parallel over the course of the experiment. Images were acquired every 10 min for a total duration of 48h using NIS software. Focus was maintained using the Nikon Perfect Focus System. A physiological temperature of 30°C was maintained on the chip using a custom sample holder with thermoelectric modules, an objective heater with heating resistors, and a PID controller (5C7-195, Oven Industries). After acquisition, NIS raw data were analyzed using the custom MATLAB software phylocell and autotrack, available on https://www.github.com/gcharvin. Cell contours and fluorescent aggregates were segmented using a modified watershed algorithm and tracking was achieved with the Hungarian method, as previously described (Goulev et al., 2017, Morlot et al., 2019). ‘Mean fluorescence per cell’ corresponds to the average GFP intensity within the cell contours. ‘Intensity after aggregation’ corresponds to the total GFP fluorescence within the segmented aggregate at the first frame where aggregation starts. ‘Intensity before aggregation’ is the total fluorescence within the cell contour in the frame preceding aggregation.

To analyze Q97-GFP and mitochondrial morphology in mammalian cells, fluorescence microscopy was performed using a Zeiss ELYRA PS.1 Super-Resolution Structured Illumination Microscope (SR-SIM) with a 63× oil immersion objective. All channels were acquired independent and subsequently. Single cell analysis was performed in this microscope as well, quantifying only the Htt-Q97 (+/− MIA40-HA) transfected cells. Among the transfected cells, those showing nuclear or cytoplasmic huntingtin aggregates were included in the category “transfected cell with aggregates”. The percentage of cells with aggregates was calculated over the total number of transfected cells. Raw confocal SIM images were processed and generated using the Black ZEN and Blue ZEN software.

## Notes

### Competing Interest Statement

The authors have declared no competing interest.

## References

Becker J, Walter W, Yan W, Craig EA (1996) Functional interaction of cytosolic hsp70 and a DnaJ-related protein, Ydj1p, in protein translocation in vivo. Mol Cell Biol 16: 4378–86

Böckler S, Chelius X, Hock N, Klecker T, Wolter M, Weiss M, Braun RJ, Westermann B (2017) Fusion, fission, and transport control asymmetric inheritance of mitochondria and protein aggregates. J Cell Biol 216: 2481–2498

Boos F, Krämer L, Groh C, Jung F, Haberkant P, Stein F, Wollweber F, Gackstatter A, Zoller E, van der Laan M, Savitski MM, Benes V, Herrmann JM (2019) Mitochondrial protein-induced stress triggers a global adaptive transcriptional programme. Nat Cell Biol 21: 442–451

Boos F, Labbadia J, Herrmann JM (2020) How the Mitoprotein-Induced Stress Response Safeguards the Cytosol: A Unified View. Trends Cell Biol 30: 241–254

Braun RJ, Buttner S, Ring J, Kroemer G, Madeo F (2010) Nervous yeast: modeling neurotoxic cell death. Trends Biochem Sci 35: 135–44

Cenini G, Rub C, Bruderek M, Voos W (2016) Amyloid beta-peptides interfere with mitochondrial preprotein import competence by a coaggregation process. Mol Biol Cell 27: 3257–3272

Chacinska A, Lind M, Frazier AE, Dudek J, Meisinger C, Geissler A, Sickmann A, Meyer HE, Truscott KN, Guiard B, Pfanner N, Rehling P (2005) Mitochondrial presequence translocase: switching between TOM tethering and motor recruitment involves Tim21 and Tim17. Cell 120: 817–29

Chacinska A, Pfannschmidt S, Wiedemann N, Kozjak V, Sanjuan Szklarz LK, Schulze-Specking A, Truscott KN, Guiard B, Meisinger C, Pfanner N (2004) Essential role of Mia40 in import and assembly of mitochondrial intermembrane space proteins. EMBO J 23: 3735–3746

Chiti F, Dobson CM (2017) Protein Misfolding, Amyloid Formation, and Human Disease: A Summary of Progress Over the Last Decade. Annu Rev Biochem 86: 27–68

Dehay B, Bertolotti A (2006) Critical role of the proline-rich region in Huntingtin for aggregation and cytotoxicity in yeast. J Biol Chem 281: 35608–15

Deshaies RJ, Koch BD, Werner-Washburne M, Craig EA, Schekman R (1988) A subfamily of stress proteins facilitates translocation of secretory and mitochondrial precursor polypeptides. Nature 332: 800–805

Douglas PM, Summers DW, Ren HY, Cyr DM (2009) Reciprocal efficiency of RNQ1 and polyglutamine detoxification in the cytosol and nucleus. Mol Biol Cell 20: 4162–73

Duennwald ML, Jagadish S, Giorgini F, Muchowski PJ, Lindquist S (2006) A network of protein interactions determines polyglutamine toxicity. Proc Natl Acad Sci U S A 103: 11051–6

Fessler E, Eckl EM, Schmitt S, Mancilla IA, Meyer-Bender MF, Hanf M, Philippou-Massier J, Krebs S, Zischka H, Jae LT (2020) A pathway coordinated by DELE1 relays mitochondrial stress to the cytosol. Nature 579: 433–437

Fischer M, Horn S, Belkacemi A, Kojer K, Petrungaro C, Habich M, Ali M, Kuttner V, Bien M, Kauff F, Dengjel J, Herrmann JM, Riemer J (2013) Protein import and oxidative folding in the mitochondrial intermembrane space of intact mammalian cells. Mol Biol Cell 24: 2160–70

Franken H, Mathieson T, Childs D, Sweetman GM, Werner T, Togel I, Doce C, Gade S, Bantscheff M, Drewes G, Reinhard FB, Huber W, Savitski MM (2015) Thermal proteome profiling for unbiased identification of direct and indirect drug targets using multiplexed quantitative mass spectrometry. Nat Protoc 10: 1567–93

Fünfschilling U, Rospert S (1999) Nascent polypeptide-associated complex stimulates protein import into yeast mitochondria. Mol Biol Cell 10: 3289–3299

Gatto L, Lilley KS (2012) MSnbase-an R/Bioconductor package for isobaric tagged mass spectrometry data visualization, processing and quantitation. Bioinformatics 28: 288–9

Gidalevitz T, Ben-Zvi A, Ho KH, Brignull HR, Morimoto RI (2006) Progressive disruption of cellular protein folding in models of polyglutamine diseases. Science 311: 1471–4

Goulev Y, Morlot S, Matifas A, Huang B, Molin M, Toledano MB, Charvin G (2017) Nonlinear feedback drives homeostatic plasticity in H2O2 stress response. Elife 6

Gruber A, Hornburg D, Antonin M, Krahmer N, Collado J, Schaffer M, Zubaite G, Luchtenborg C, Sachsenheimer T, Brugger B, Mann M, Baumeister W, Hartl FU, Hipp MS, Fernandez-Busnadiego R (2018) Molecular and structural architecture of polyQ aggregates in yeast. Proc Natl Acad Sci U S A

Guo PC, Ma JD, Jiang YL, Wang SJ, Bao ZZ, Yu XJ, Chen Y, Zhou CZ (2012) Structure of yeast sulfhydryl oxidase erv1 reveals electron transfer of the disulfide relay system in the mitochondrial intermembrane space. J Biol Chem 287: 34961–9

Guo X, Aviles G, Liu Y, Tian R, Unger BA, Lin YT, Wiita AP, Xu K, Correia MA, Kampmann M (2020) Mitochondrial stress is relayed to the cytosol by an OMA1-DELE1-HRI pathway. Nature 579: 427–432

Guo X, Sun X, Hu D, Wang YJ, Fujioka H, Vyas R, Chakrapani S, Joshi AU, Luo Y, Mochly-Rosen D, Qi X (2016) VCP recruitment to mitochondria causes mitophagy impairment and neurodegeneration in models of Huntington’s disease. Nat Commun 7: 12646

Habich M, Salscheider SL, Murschall LM, Hoehne MN, Fischer M, Schorn F, Petrungaro C, Ali M, Erdogan AJ, Abou-Eid S, Kashkar H, Dengjel J, Riemer J (2019) Vectorial Import via a Metastable Disulfide-Linked Complex Allows for a Quality Control Step and Import by the Mitochondrial Disulfide Relay. Cell Rep 26: 759–774 e5

Hangen E, Feraud O, Lachkar S, Mou H, Doti N, Fimia GM, Lam NV, Zhu C, Godin I, Muller K, Chatzi A, Nuebel E, Ciccosanti F, Flamant S, Benit P, Perfettini JL, Sauvat A, Bennaceur-Griscelli A, Ser-Le Roux K, Gonin P et al. (2015) Interaction between AIF and CHCHD4 Regulates Respiratory Chain Biogenesis. Mol Cell 58: 1001–14

Hansen KG, Aviram N, Laborenz J, Bibi C, Meyer M, Spang A, Schuldiner M, Herrmann JM (2018) An ER surface retrieval pathway safeguards the import of mitochondrial membrane proteins in yeast. Science 361: 1118–1122

Hibbs MA, Hess DC, Myers CL, Huttenhower C, Li K, Troyanskaya OG (2007) Exploring the functional landscape of gene expression: directed search of large microarray compendia. Bioinformatics 23: 2692–9

Higgins R, Kabbaj MH, Hatcher A, Wang Y (2018) The absence of specific yeast heat-shock proteins leads to abnormal aggregation and compromised autophagic clearance of mutant Huntingtin proteins. PLoS One 13: e0191490

Higurashi T, Hines JK, Sahi C, Aron R, Craig EA (2008) Specificity of the J-protein Sis1 in the propagation of 3 yeast prions. Proc Natl Acad Sci U S A 105: 16596–601

Hoseini H, Pandey S, Jores T, Schmitt A, Franz-Wachtel M, Macek B, Buchner J, Dimmer KS, Rapaport D (2016) The cytosolic cochaperone Sti1 is relevant for mitochondrial biogenesis and morphology. FEBS J 283: 3338–52

Hoshino A, Wang WJ, Wada S, McDermott-Roe C, Evans CS, Gosis B, Morley MP, Rathi KS, Li J, Li K, Yang S, McManus MJ, Bowman C, Potluri P, Levin M, Damrauer S, Wallace DC, Holzbaur ELF, Arany Z (2019) The ADP/ATP translocase drives mitophagy independent of nucleotide exchange. Nature 575: 375–379

Huber W, von Heydebreck A, Sultmann H, Poustka A, Vingron M (2002) Variance stabilization applied to microarray data calibration and to the quantification of differential expression. Bioinformatics 18 Suppl 1: S96–104

Hughes CS, Foehr S, Garfield DA, Furlong EE, Steinmetz LM, Krijgsveld J (2014) Ultrasensitive proteome analysis using paramagnetic bead technology. Mol Syst Biol 10: 757

Hwang S, Disatnik MH, Mochly-Rosen D (2015) Impaired GAPDH-induced mitophagy contributes to the pathology of Huntington’s disease. EMBO Mol Med 7: 1307–26

Jores T, Lawatscheck J, Beke V, Franz-Wachtel M, Yunoki K, Fitzgerald JC, Macek B, Endo T, Kalbacher H, Buchner J, Rapaport D (2018) Cytosolic Hsp70 and Hsp40 chaperones enable the biogenesis of mitochondrial beta-barrel proteins. J Cell Biol 217: 3091–3108

Khalil B, El Fissi N, Aouane A, Cabirol-Pol MJ, Rival T, Lievens JC (2015) PINK1-induced mitophagy promotes neuroprotection in Huntington’s disease. Cell Death Dis 6: e1617

Kim YE, Hosp F, Frottin F, Ge H, Mann M, Hayer-Hartl M, Hartl FU (2016) Soluble Oligomers of PolyQ-Expanded Huntingtin Target a Multiplicity of Key Cellular Factors. Mol Cell 63: 951–64

Klaips CL, Gropp MHM, Hipp MS, Hartl FU (2020) Sis1 potentiates the stress response to protein aggregation and elevated temperature. Nat Commun 11: 6271

Koehler CM, Jarosch E, Tokatlidis K, Schmid K, Schweyen RJ, Schatz G (1998) Import of mitochondrial carrier proteins mediated by essential proteins of the intermembrane space. Science 279: 369–373

Kowalski L, Bragoszewski P, Khmelinskii A, Glow E, Knop M, Chacinska A (2018) Determinants of the cytosolic turnover of mitochondrial intermembrane space proteins. BMC Biol 16: 66

Krämer L, Groh C, Herrmann JM (2020) The proteasome: friend and foe of mitochondrial biogenesis. FEBS Lett

Krobitsch S, Lindquist S (2000) Aggregation of huntingtin in yeast varies with the length of the polyglutamine expansion and the expression of chaperone proteins. Proc Natl Acad Sci U S A 97: 1589–94

Labbadia J, Brielmann RM, Neto MF, Lin YF, Haynes CM, Morimoto RI (2017) Mitochondrial Stress Restores the Heat Shock Response and Prevents Proteostasis Collapse during Aging. Cell Rep 21: 1481–1494

Laborenz J, Hansen K, Prescianotto-Baschong C, Spang A, Herrmann JM (2019) In vitro import experiments with semi-intact cells suggest a role of the Sec61 paralog Ssh1 in mitochondrial biogenesis. Biol Chem 400: 1229–1240

Lehmer C, Schludi MH, Ransom L, Greiling J, Junghanel M, Exner N, Riemenschneider H, van der Zee J, Van Broeckhoven C, Weydt P, Heneka MT, Edbauer D (2018) A novel CHCHD10 mutation implicates a Mia40-dependent mitochondrial import deficit in ALS. EMBO Mol Med 10

Li Q, Vande Velde C, Israelson A, Xie J, Bailey AO, Dong MQ, Chun SJ, Roy T, Winer L, Yates JR, Capaldi RA, Cleveland DW, Miller TM (2010) ALS-linked mutant superoxide dismutase 1 (SOD1) alters mitochondrial protein composition and decreases protein import. Proc Natl Acad Sci U S A 107: 21146–51

Li Y, Xue Y, Xu X, Wang G, Liu Y, Wu H, Li W, Wang Y, Chen Z, Zhang W, Zhu Y, Ji W, Xu T, Liu L, Chen Q (2019) A mitochondrial FUNDC1/HSC70 interaction organizes the proteostatic stress response at the risk of cell morbidity. EMBO J 38

Liu Y, Wang X, Coyne LP, Yang Y, Qi Y, Middleton FA, Chen XJ (2019) Mitochondrial carrier protein overloading and misfolding induce aggresomes and proteostatic adaptations in the cytosol. Mol Biol Cell 30: 1272–1284

Lu K, Psakhye I, Jentsch S (2014) Autophagic clearance of polyQ proteins mediated by ubiquitin-Atg8 adaptors of the conserved CUET protein family. Cell 158: 549–63

Luciano P, Vial S, Vergnolle MA, Dyall SD, Robinson DR, Tokatlidis K (2001) Functional reconstitution of the import of the yeast ADP/ATP carrier mediated by the TIM10 complex. EMBO J 20: 4099–4106

Mårtensson CU, Priesnitz C, Song J, Ellenrieder L, Doan KN, Boos F, Floerchinger A, Zufall N, Oeljeklaus S, Warscheid B, Becker T (2019) Mitochondrial protein translocation-associated degradation. Nature 569: 679–683

Mason RP, Casu M, Butler N, Breda C, Campesan S, Clapp J, Green EW, Dhulkhed D, Kyriacou CP, Giorgini F (2013) Glutathione peroxidase activity is neuroprotective in models of Huntington’s disease. Nat Genet 45: 1249–54

Meriin AB, Zhang X, He X, Newnam GP, Chernoff YO, Sherman MY (2002) Huntington toxicity in yeast model depends on polyglutamine aggregation mediated by a prion-like protein Rnq1. J Cell Biol 157: 997–1004

Metzger MB, Michaelis S (2009) Analysis of quality control substrates in distinct cellular compartments reveals a unique role for Rpn4p in tolerating misfolded membrane proteins. Mol Biol Cell 20: 1006–19

Moggridge S, Sorensen PH, Morin GB, Hughes CS (2018) Extending the Compatibility of the SP3 Paramagnetic Bead Processing Approach for Proteomics. J Proteome Res 17: 1730–1740

Mogk A, Bukau B (2014) Mitochondria tether protein trash to rejuvenate cellular environments. Cell 159: 471–2

Mohanraj K, Wasilewski M, Beninca C, Cysewski D, Poznanski J, Sakowska P, Bugajska Z, Deckers M, Dennerlein S, Fernandez-Vizarra E, Rehling P, Dadlez M, Zeviani M, Chacinska A (2019) Inhibition of proteasome rescues a pathogenic variant of respiratory chain assembly factor COA7. EMBO Mol Med 11: e9561

Morlot S, Song J, Leger-Silvestre I, Matifas A, Gadal O, Charvin G (2019) Excessive rDNA Transcription Drives the Disruption in Nuclear Homeostasis during Entry into Senescence in Budding Yeast. Cell Rep 28: 408–422 e4

Mossmann D, Vogtle FN, Taskin AA, Teixeira PF, Ring J, Burkhart JM, Burger N, Pinho CM, Tadic J, Loreth D, Graff C, Metzger F, Sickmann A, Kretz O, Wiedemann N, Zahedi RP, Madeo F, Glaser E, Meisinger C (2014) Amyloid-beta peptide induces mitochondrial dysfunction by inhibition of preprotein maturation. Cell Metab 20: 662–9

Murschall LM, Gerhards A, MacVicar T, Peker E, Hasberg L, Wawra S, Langer T, Riemer J (2020) The C-terminal region of the oxidoreductase MIA40 stabilizes its cytosolic precursor during mitochondrial import. BMC Biol 18: 96

Naoe M, Ohwa Y, Ishikawa D, Ohshima C, Nishikawa S, Yamamoto H, Endo T (2004) Identification of Tim40 that mediates protein sorting to the mitochondrial intermembrane space. J Biol Chem 279: 47815–47821

Napoli E, Wong S, Hung C, Ross-Inta C, Bomdica P, Giulivi C (2013) Defective mitochondrial disulfide relay system, altered mitochondrial morphology and function in Huntington’s disease. Hum Mol Genet 22: 989–1004

Opalinski L, Song J, Priesnitz C, Wenz LS, Oeljeklaus S, Warscheid B, Pfanner N, Becker T (2018) Recruitment of Cytosolic J-Proteins by TOM Receptors Promotes Mitochondrial Protein Biogenesis. Cell Rep 25: 2036–2043 e5

Papic D, Elbaz-Alon Y, Koerdt SN, Leopold K, Worm D, Jung M, Schuldiner M, Rapaport D (2013) The role of Djp1 in import of the mitochondrial protein Mim1 demonstrates specificity between a cochaperone and its substrate protein. Mol Cell Biol 33: 4083–94

Papsdorf K, Kaiser CJ, Drazic A, Grotzinger SW, Haessner C, Eisenreich W, Richter K (2015) Polyglutamine toxicity in yeast induces metabolic alterations and mitochondrial defects. BMC Genomics 16: 662

Peleh V, Cordat E, Herrmann JM (2016) Mia40 is a trans-site receptor that drives protein import into the mitochondrial intermembrane space by hydrophobic substrate binding. Elife 5: e16177

Peleh V, Zannini F, Backes S, Rouhier N, Herrmann JM (2017) Erv1 of Arabidopsis thaliana can directly oxidize mitochondrial intermembrane space proteins in the absence of redox-active Mia40. BMC Biol 15: 106

Perez-Riverol Y, Csordas A, Bai J, Bernal-Llinares M, Hewapathirana S, Kundu DJ, Inuganti A, Griss J, Mayer G, Eisenacher M, Perez E, Uszkoreit J, Pfeuffer J, Sachsenberg T, Yilmaz S, Tiwary S, Cox J, Audain E, Walzer M, Jarnuczak AF et al. (2019) The PRIDE database and related tools and resources in 2019: improving support for quantification data. Nucleic Acids Res 47: D442–D450

Piard J, Umanah GKE, Harms FL, Abalde-Atristain L, Amram D, Chang M, Chen R, Alawi M, Salpietro V, Rees MI, Chung SK, Houlden H, Verloes A, Dawson TM, Dawson VL, Van Maldergem L, Kutsche K (2018) A homozygous ATAD1 mutation impairs postsynaptic AMPA receptor trafficking and causes a lethal encephalopathy. Brain 141: 651–661

Pihlasalo S, Kirjavainen J, Hanninen P, Harma H (2011) High sensitivity luminescence nanoparticle assay for the detection of protein aggregation. Anal Chem 83: 1163–6

Ritchie ME, Phipson B, Wu D, Hu Y, Law CW, Shi W, Smyth GK (2015) limma powers differential expression analyses for RNA-sequencing and microarray studies. Nucleic Acids Res 43: e47

Ruan L, Zhou C, Jin E, Kucharavy A, Zhang Y, Wen Z, Florens L, Li R (2017) Cytosolic proteostasis through importing of misfolded proteins into mitochondria. Nature 543: 443–446

Saladi S, Boos F, Poglitsch M, Meyer H, Sommer F, Muhlhaus T, Schroda M, Schuldiner M, Madeo F, Herrmann JM (2020) The NADH Dehydrogenase Nde1 Executes Cell Death after Integrating Signals from Metabolism and Proteostasis on the Mitochondrial Surface. Mol Cell 77: 189–202 e6

Savitski MM, Wilhelm M, Hahne H, Kuster B, Bantscheff M (2015) A Scalable Approach for Protein False Discovery Rate Estimation in Large Proteomic Data Sets. Mol Cell Proteomics 14: 2394–404

Shakya VPS, Barbeau WA, Xiao T, Knutson CS, Hughes AL (2020) The nucleus is a quality control center for non-imported mitochondrial proteins. bioRxiv: 2020.06.26.173781

Sideri TC, Koloteva-Levine N, Tuite MF, Grant CM (2011) Methionine oxidation of Sup35 protein induces formation of the [PSI+] prion in a yeast peroxiredoxin mutant. J Biol Chem 286: 38924–31

Solans A, Zambrano A, Rodriguez M, Barrientos A (2006) Cytotoxicity of a mutant huntingtin fragment in yeast involves early alterations in mitochondrial OXPHOS complexes II and III. Hum Mol Genet 15: 3063–81

Sontag EM, Samant RS, Frydman J (2017) Mechanisms and Functions of Spatial Protein Quality Control. Annu Rev Biochem 86: 97–122

Sorrentino V, Romani M, Mouchiroud L, Beck JS, Zhang H, D’Amico D, Moullan N, Potenza F, Schmid AW, Rietsch S, Counts SE, Auwerx J (2017) Enhancing mitochondrial proteostasis reduces amyloid-beta proteotoxicity. Nature 552: 187–193

Sridharan S, Kurzawa N, Werner T, Gunthner I, Helm D, Huber W, Bantscheff M, Savitski MM (2019) Proteome-wide solubility and thermal stability profiling reveals distinct regulatory roles for ATP. Nat Commun 10: 1155

Strimmer K (2008) fdrtool: a versatile R package for estimating local and tail area-based false discovery rates. Bioinformatics 24: 1461–2

Terada K, Kanazawa M, Bukau B, Mori M (1997) The human DnaJ homologue dj2 facilitates mitochondrial protein import and luciferase refolding. J Cell Biol 139: 1089–95

Terziyska N, Grumbt B, Bien M, Neupert W, Herrmann JM, Hel K (2007) The sulfhydryl oxidase Erv1 is a substrate of the Mia40-dependent protein translocation pathway. FEBS Lett 581: 1098–1102

Terziyska N, Lutz T, Kozany C, Mokranjac D, Mesecke N, Neupert W, Herrmann JM, Hell K (2005) Mia40, a novel factor for protein import into the intermembrane space of mitochondria is able to bind metal ions. FEBS Lett 579: 179–184

van Well EM, Bader V, Patra M, Sanchez-Vicente A, Meschede J, Furthmann N, Schnack C, Blusch A, Longworth J, Petrasch-Parwez E, Mori K, Arzberger T, Trumbach D, Angersbach L, Showkat C, Sehr DA, Berlemann LA, Goldmann P, Clement AM, Behl C et al. (2019) A protein quality control pathway regulated by linear ubiquitination. EMBO J 38

Vaquer-Alicea J, Diamond MI (2019) Propagation of Protein Aggregation in Neurodegenerative Diseases. Annu Rev Biochem 88: 785–810

Vartiainen S, Chen S, George J, Tuomela T, Luoto KR, O’Dell KM, Jacobs HT (2014) Phenotypic rescue of a Drosophila model of mitochondrial ANT1 disease. Dis Model Mech 7: 635–48

Wang X, Chen XJ (2015) A cytosolic network suppressing mitochondria-mediated proteostatic stress and cell death. Nature 524: 481–4

Weidberg H, Amon A (2018) MitoCPR-A surveillance pathway that protects mitochondria in response to protein import stress. Science 360: eaan4146

Werner T, Sweetman G, Savitski MF, Mathieson T, Bantscheff M, Savitski MM (2014) Ion coalescence of neutron encoded TMT 10-plex reporter ions. Anal Chem 86: 3594–601

Williams CC, Jan CH, Weissman JS (2014) Targeting and plasticity of mitochondrial proteins revealed by proximity-specific ribosome profiling. Science 346: 748–51

Wolfe KJ, Ren HY, Trepte P, Cyr DM (2013) The Hsp70/90 cochaperone, Sti1, suppresses proteotoxicity by regulating spatial quality control of amyloid-like proteins. Mol Biol Cell 24: 3588–602

Wrobel L, Topf U, Bragoszewski P, Wiese S, Sztolsztener ME, Oeljeklaus S, Varabyova A, Lirski M, Chroscicki P, Mroczek S, Januszewicz E, Dziembowski A, Koblowska M, Warscheid B, Chacinska A (2015) Mistargeted mitochondrial proteins activate a proteostatic response in the cytosol. Nature 524: 485–8

Wu Z, Senchuk MM, Dues DJ, Johnson BK, Cooper JF, Lew L, Machiela E, Schaar CE, DeJonge H, Blackwell TK, Van Raamsdonk JM (2018) Mitochondrial unfolded protein response transcription factor ATFS-1 promotes longevity in a long-lived mitochondrial mutant through activation of stress response pathways. BMC Biol 16: 147

Xie Y, Varshavsky A (2001) RPN4 is a ligand, substrate, and transcriptional regulator of the 26S proteasome: a negative feedback circuit. Proc Natl Acad Sci U S A 98: 3056–61

Yablonska S, Ganesan V, Ferrando LM, Kim J, Pyzel A, Baranova OV, Khattar NK, Larkin TM, Baranov SV, Chen N, Strohlein CE, Stevens DA, Wang X, Chang YF, Schurdak ME, Carlisle DL, Minden JS, Friedlander RM (2019) Mutant huntingtin disrupts mitochondrial proteostasis by interacting with TIM23. Proc Natl Acad Sci U S A 116: 16593–16602

Yang J, Hao X, Cao X, Liu B, Nystrom T (2016) Spatial sequestration and detoxification of Huntingtin by the ribosome quality control complex. Elife 5

Yano H, Baranov SV, Baranova OV, Kim J, Pan Y, Yablonska S, Carlisle DL, Ferrante RJ, Kim AH, Friedlander RM (2014) Inhibition of mitochondrial protein import by mutant huntingtin. Nat Neurosci 17: 822–31

Zhang Q, Wu X, Chen P, Liu L, Xin N, Tian Y, Dillin A (2018) The Mitochondrial Unfolded Protein Response Is Mediated Cell-Non-autonomously by Retromer-Dependent Wnt Signaling. Cell 174: 870–883 e17

Zheng X, Krakowiak J, Patel N, Beyzavi A, Ezike J, Khalil AS, Pincus D (2016) Dynamic control of Hsf1 during heat shock by a chaperone switch and phosphorylation. Elife 5

Zhou C, Slaughter BD, Unruh JR, Eldakak A, Rubinstein B, Li R (2011) Motility and segregation of Hsp104-associated protein aggregates in budding yeast. Cell 147: 1186–96

Zöller E, Laborenz J, Kramer L, Boos F, Raschle M, Alexander RT, Herrmann JM (2020) The intermembrane space protein Mix23 is a novel stress-induced mitochondrial import factor. J Biol Chem 295: 14686–14697

